# Single-cell transcriptional profiling of clear cell renal cell carcinoma reveals an invasive tumor vasculature phenotype

**DOI:** 10.1101/2023.08.09.552568

**Authors:** Justina Zvirblyte, Juozas Nainys, Simonas Juzenas, Raimonda Kubiliute, Marius Kincius, Albertas Ulys, Sonata Jarmalaite, Linas Mazutis

## Abstract

Clear cell renal cell carcinoma (ccRCC) is the most prevalent form of renal cancer, accounting for over 75% of cases. The asymptomatic nature of the disease contributes to late-stage diagnoses and poor survival. Highly vascularized and immune infiltrated microenvironment are prominent features of ccRCC, yet the interplay between vasculature and immune cells, disease progression and response to therapy remains poorly understood. Using droplet-based single-cell RNA sequencing we profiled 50,236 transcriptomes from paired tumor and healthy adjacent kidney tissues. Our analysis revealed significant heterogeneity and inter-patient variability of the tumor microenvironment. Notably, we discovered a previously uncharacterized vasculature subpopulation associated with epithelial-mesenchymal transition. The cell-cell communication analysis revealed multiple modes of immunosuppressive interactions within the tumor microenvironment, including clinically relevant interactions between tumor vasculature and stromal cells with immune cells. The upregulation of the genes involved in these interactions was associated with worse survival in the TCGA KIRC cohort. Our findings demonstrate the role of tumor vasculature and stromal cell populations in shaping the ccRCC microenvironment and uncover a subpopulation of cells within the tumor vasculature that is associated with an invasive phenotype.

## INTRODUCTION

The asymptomatic nature of clear cell renal cell carcinoma (ccRCC), the most common renal cancer, often leads to diagnosis in late III or IV stage with survival probability of 59% and 20%, respectively. Approximately 30% of cases metastasize^1^. Previous efforts aimed at characterizing ccRCC tumors have provided valuable insights into the genomic^2^, transcriptomic and epigenetic^3,4^ landscape of both the tumor and the tumor microenvironment (TME). It is now well-established that the most abundant genomic alterations in ccRCC involve the loss of regions in 3p chromosome (occurring in >90% of cases) and von Hippel–Lindau (*VHL*) gene mutations (>50% of cases). These alterations lead to impaired degradation and abnormal accumulation of hypoxia-inducible factors (HIFs)^2,3^, resulting in a highly vascularized tumor appearance. Moreover, ccRCC tumors exhibit a high degree of immune infiltration^5,6^. Consequently, the most common first-line treatment options for the localized disease involve surgical removal of the tumor, while advanced disease may be treated with VEGF pathway inhibitors, standalone or in combination with immune checkpoint blockade therapies^2,7,8^. However, owing to a high degree of intra- and inter-tumor heterogeneity, these treatments benefit only a fraction of patients, and often result in acquired resistance and further disease progression^2,9^.

Recent advancements in microfluidics and molecular barcoding have enabled high-throughput transcriptional, epigenomic and even multi-omic tissue profiling at the single cell resolution, yielding important biological insights. For instance, using single-cell RNA sequencing (scRNA-Seq) a plethora of single-cell resolution healthy and cancerous tissue atlases have been constructed, revealing the phenotypic complexity and plasticity of the tumor microenvironment^10–13^. In the context of ccRCC, single-cell techniques have shed light on the cell of origin of ccRCC^14,15^, malignancy-related transcriptional programs of the tumor^16^ and the heterogeneous tumor-associated immune cell infiltrate^17–20^. Furthermore, the phenotypical changes of immune cell populations along advancing disease stage^21^ and immunotherapy treatment^18,22^ have been characterized in detail.

Upon the widespread adoption of the single cell profiling techniques there was a noticeable paradigm shift in the field of cancer research – a systemic view of the tumor as a highly orchestrated ecosystem took over the tumor cell-centric point of view. This shift has highlighted the crucial role of other players in the TME, including various subpopulations of stromal and endothelial cells that have been discovered to have an impact on disease progression, response to therapy and patient survival^23,24^. While considerable efforts have been made to characterize the ccRCC tumor microenvironment at the single cell level, most of the previous studies focused on tumor or immune cells, leaving the role of other cells types within the ccRCC TME poorly understood. In this study, we aimed to address this gap by profiling fresh ccRCC tumor and matched healthy adjacent tissue samples using droplet-based scRNA-Seq, omitting cell sorting and enrichment steps in order to capture the diverse phenotypes present in the TME, including the stromal cell populations. As a result, we captured all major specialized epithelial and endothelial cell populations in healthy adjacent kidney tissue, including a progenitor-like epithelial cell phenotype resembling the cell of origin for ccRCC. Furthermore, we described five tumor endothelium subpopulations and discovered a previously uncharacterized tip-like cell phenotype. Within the TME, we identifed well-described immunosuppressive tumor associated macrophage (TAM) populations and exhausted infiltrating T cells^21^. Through cell-cell communication analysis, we inferred the interactions between various cell types within the TME, revealing tumor vasculature and stromal cell involvement in maintaining an immunosuppressive niche. Expression of genes involved in these interactions was associated with worse overall survival in the TCGA KIRC cohort. Overall, our results complement ongoing ccRCC TME characterization efforts by introducing a novel endothelial phenotype and highlighting the importance as well as potential therapeutic relevance of stromal and endothelial cells in the TME.

## RESULTS

### Single cell profiling of healthy and tumor tissues reveals inter-patient variability and epithelial ccRCC progenitor-like population in healthy tissue

To dissect the transcriptional landscape of the human ccRCC tumor microenvironment (TME), we profiled fresh tumor (n=8) and healthy adjacent (n=9) kidney tissue samples using a droplet-based scRNA-seq platform (Figure 1a). To capture the diverse range of cell types constituting the TME, our experimental strategy involved rapid isolation of dissociated cells in microfluidic droplets, without any enrichment or sorting steps (see Methods). Following quality control, batch correction and doublet removal (see Methods), we obtained a total of 50,236 single cell transcriptomes that were then clustered using a graph-based spectral clustering. The cell types belonging to each cluster were identified manually based on differentially expressed top 25 marker genes (adjusted p-value <0.05; cluster vs the rest of cells, Mann-Whitney U test with Benjamini-Hochberg correction), validated by extensive literature review (Figure 1b, f and Supplementary file Table 1).

**Figure 1.**
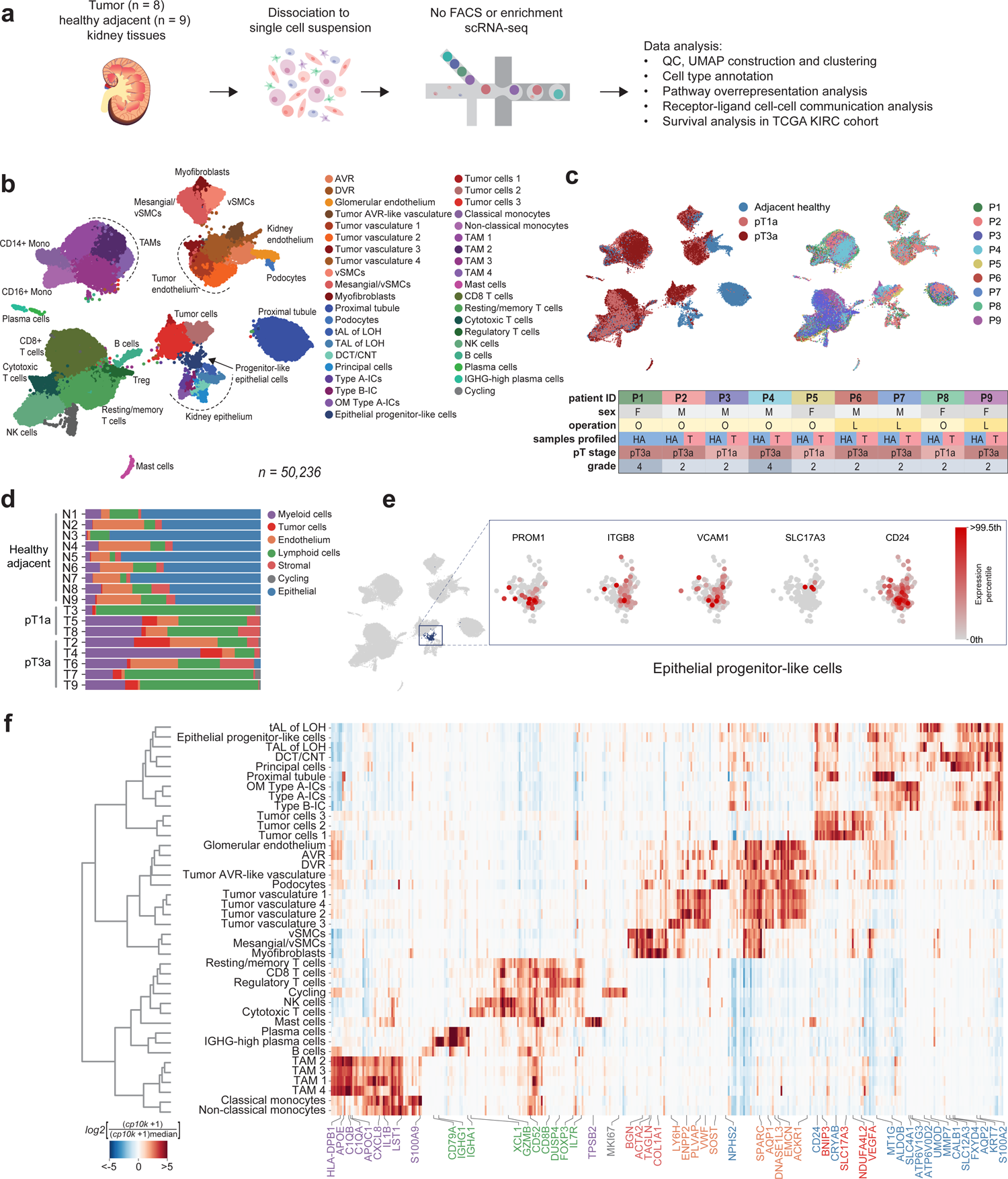
Profiling the ccRCC microenvironment. a) Experimental design. b) Global single cell transcriptional map of ccRCC. c) Clinical information of collected samples and corresponding UMAPs of cells annotated by disease stage (adjacent healthy, pT1a and pT3a) and patient ID (P1-P9). Healthy adjacent samples (blue) almost completely separate from the tumor (light and dark red). d) Sample composition by major cell type. Notably, healthy adjacent samples are enriched with specialized kidney epithelial and endothelial cells, while tumor samples are enriched for immune cells. e) Expression of ccRCC cell of origin markers in epithelial progenitor-like cell population. f) Global heatmap for population specific markers. Only genes with Benjamini-Hochberg adjusted p-value <0.05 are shown. Color of the gene name indicates major cell type. AVR – ascending vasa recta, DVR – descending vasa recta, vSMCs – vascular smooth muscle cells, LOH – loop of Henle, tAL – thin ascending limb, TAL – thick ascending limb, DCT/CNT – distal convoluted/connecting tubule, ICs – intercalated cells, OM – outer medullary, TAM – tumor associated macrophages.

**Table 1.** Reagents and materials used in the study.

| Resource | Source | Identifier/<br>Cat No. |
| --- | --- | --- |
| ccRCC and paired healthy adjacent kidney samples | National Cancer Institute, Vilnius, Lithuania | N/A |
| Tumor Dissociation Kit | Miltenyi Biotec | 130-095-929 |
| Tissue Dissociation Kit I | Miltenyi Biotec | 130-110-201 |
| RBC lysis reagent | Miltenyi Biotec | 130-094-183 |
| DPBS | Gibco | 14080-048 |
| Trypan Blue solution | Gibco | 15250061 |
| Maxima H- minus reverse transcriptase | Thermo Scientific | EP0751 |
| dNTP (10 mM each) | Thermo Scientific | R0192 |
| RiboLock RNase inhibitor | Thermo Scientific | EO0382 |
| Igepal CA-630 | Sigma Aldrich | 18896-50mL |
| AMPure XP reagent | BeckMan Coulter | A63881 |
| 2X KAPA HiFi Hot Start Ready Mix | Roche | KK2601 |
| NEBNext Ultra II FS DNA Library Prep Kit | NEB | E7805S |
| Bioanalyzer DNA High Sensitivity assay | Agilent | 50674626 |
| NextSeq 500/550 HO Kit v2.5 (75 Cycles) | Illumina | 20024906 |
| NextSeq 500/550 HO Kit v2.5 (150 Cycles) | Illumina | 20024907 |

Healthy-adjacent samples displayed all major epithelial and endothelial cell populations characteristic of a healthy kidney (Figure 1b)^25–27^. By omitting the cell enrichment step, we could successfully capture diverse cell types that are known to be highly sensitive to handling and extended workflow procedures^28^. For example, we captured both, ascending (*DNASE1L3*) and descending (*AQP1*, *SLC14A1*) parts of the vasa recta, as well as glomerular endothelium marked by *IGFBP5* and *SOST* expression. The epithelial compartment encompassed cells from various specialized nephron segments, including rare populations such as intercalating cells of type A and B (expressing marker genes *ATP6V1G3* and *SLC26A4*, respectively), as well as podocytes (*NPHS2*, *PODXL*). Interestingly, in contrast to tumor, all healthy tissue samples comprised a population of epithelial progenitor-like cells, similar to that described by *Young et al*.^14^ (Figure 1e). This population expressed genes associated with de-differentiated injured kidney epithelium, such as *PROM1* and *ITGB8* ^29^, as well as *CD24* and *SOX4*, which have been implicated in kidney development and mark proximal tubule and distal nephron response to acute kidney injury^30^. Therefore, the epithelial progenitor-like cell population in our dataset likely represents a de-differentiated phenotype, and a potential cell of origin for ccRCC disease.

The tumor samples encompassed localized and locally advanced pT1a and pT3a pathologic stages of ccRCC (Figure 1c, Supplementary table S1). These samples exhibited high immune cell infiltration, including several populations of tumor-associated macrophages and T cells (Figure 1b). The stromal cells separated into myofibroblast (type I, IV and VI collagens, *FN1*, *TIMP2*, *ACTA2*), vascular smooth muscle cell (*TAGLN*, *ACTA2*, *SNCG*) and mesangial/vSMC (*BGN*, *PDGFRB*, *TAGLN*) clusters. Tumor endothelium completely separated from healthy-adjacent endothelial populations (Figure 1b) and included ascending vasa recta-like cells (*ACKR1*, *DNASE1L3*) as well as heterogeneous vasculature subpopulations expressing tumor-associated endothelial markers *PLVAP*, *VWF*, *SPARC*, *INSR*, *ANGPT2,* and others (Supplementary tables S2, S3). Tumor vasculature exhibited distinct expression patterns as compared to healthy endothelium (Figure 1f, 3b). While four out of five vasculature subpopulations identified in our data have been described previously^14–16^, one tumor vasculature subpopulation (Tumor vasculature 3 comprising 151 cells) appeared to be novel in the context of ccRCC and featured upregulation of *LY6H*, *PGF*, *LOX*, *CHST1* and type IV collagen (Figure 1f, 3c), consistent with a tip-cell phenotype^31^.

The tumor cells in all samples expressed canonical markers *CA9*, *NDUFA4L2*, *VEGFA* and segregated into three subpopulations, out of which one (Tumor cells 1) was patient-specific (126 cells in population, Supplementary figure S1a, b). Notably, these cells exhibited elevated expression of progenitor-like phenotype marker *SLC17A3,* which was not highly expressed in the healthy-adjacent epithelial progenitor cells (Figure 1e, Supplementary figure S1b). Furthermore, Tumor cells 1 population was the most distinct from other tumor cells based on unsupervised hierarchical clustering (Figure 1f, Supplementary figure S1b). These cells over-expressed genes such as vitamin D binding protein *GC* and *HLA-G,* the latter being involved in immunosuppressive interactions (Figure 2c), as well as *FABP7*, crucial for lipid uptake and storage in hypoxic conditions when *de novo* lipid synthesis is repressed^32^. Additionally, these cells were marked by high expression of pan-cancer marker *MDK*^33^, along with *IFI27* and *SOD2* (Supplementary figure S1b), both of which play a role in interferon response^22^. Consistently, Tumor cells 1 was the only tumor cell population not enriched for hypoxia, but instead enriched for oxidative phosphorylation and adipogenesis (Figure 4a). Considering the elevated expression of *VCAM1* and *SLC17A3*, it is possible to envision that this small patient-specific population could represent an intermediate progenitor-tumor cell phenotype.

**Figure 2.**
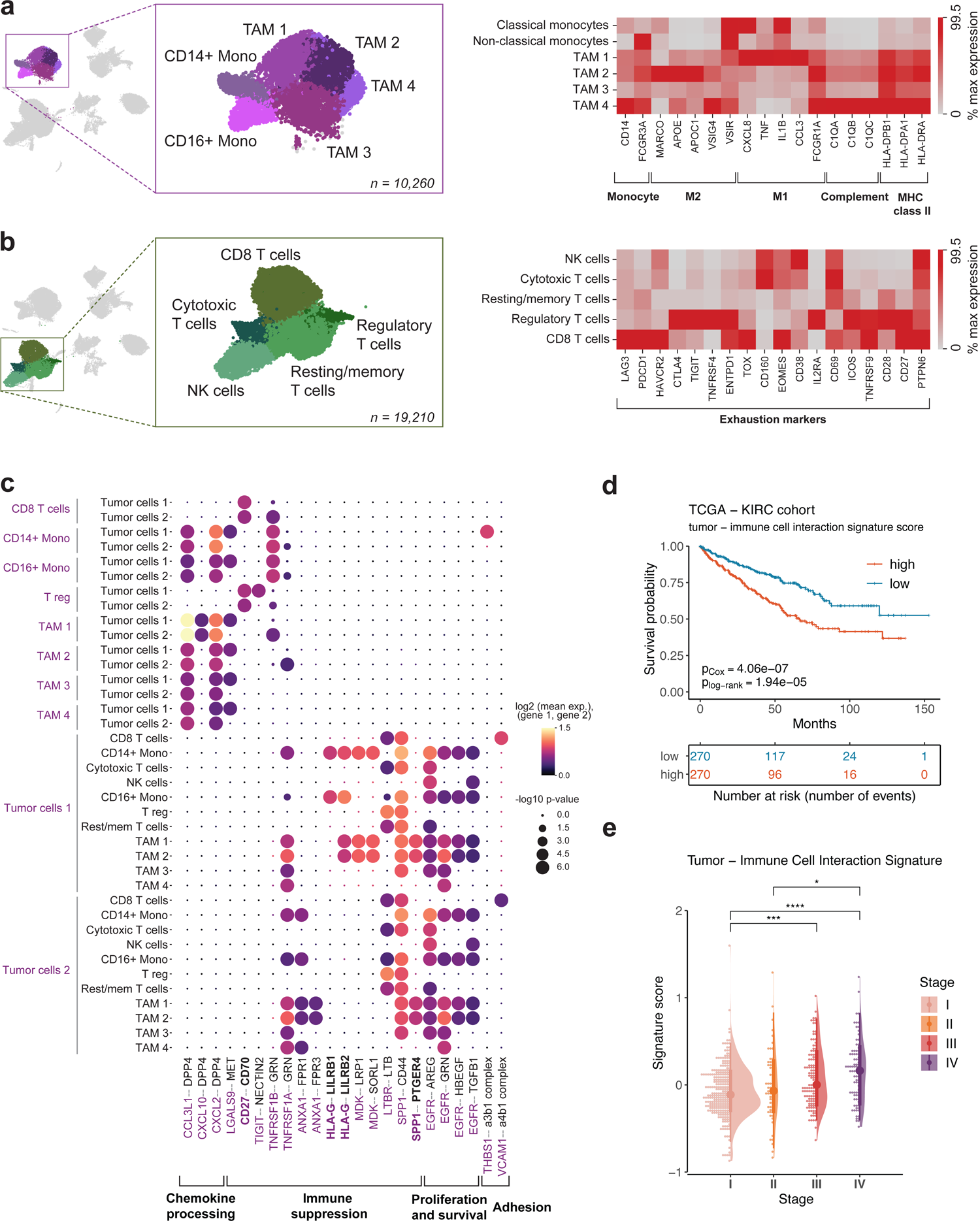
Characterization of immune cell populations found in ccRCC. a) Myeloid cell compartment consists of CD14+ and CD16+ monocytes and 4 populations of tumor associated macrophages diverse in expression of polarization markers. b) Lymphoid cells in ccRCC display heterogeneous exhaustion profile. c) Immunosuppressive interactions of clinical importance revealed by cell-cell communication analysis between immune and tumor cells using CellPhoneDB. d) Tumor-immune cell interaction signature expression in TCGA KIRC cohort is associated with a worse overall survival. e) Tumor-immune cell interaction signature increases along the progression of the ccRCC disease.

The cellular composition of tumor tissues, as expected, displayed noticeable variability across the patients as compared to their matched pair of healthy-adjacent tissues (Figure 1d, Supplementary Table S4). A common theme to all tumor samples was a high number of immune cells infiltrating the TME, accompanied by almost complete loss of specialized kidney-specific epithelial and endothelial cell populations (Figure 1c, d and Supplementary figure S1a). Except for Tumor cells 1, no other cell phenotype was patient-specific; cell population composition analysis by patient ID confirmed adequate representation of cells of different origin (Supplementary figure S1a). To quantitively assess tumor sample heterogeneity, we calculated Shannon entropy for each broad cell category^11^. Low entropy values for a cell phenotype indicate that it is rarely shared between samples, meaning that the level of heterogeneity within samples is high. In tumor samples, the heterogeneity was highest for stromal, endothelial and tumor cells, whereas healthy adjacent tissue samples exhibited comparatively lower heterogeneity (Supplementary figure S1c, d). Such diverse TME snapshots among different patients in our and other ccRCC studies^15,34^ suggest that patient stratification may rely on the abundance of specific cellular phenotypes within the TME, rather than patient-specific phenotypes. This underscores the importance of revisiting strategies for biomarker selection to aid personalized treatment options in ccRCC.

### Tumor associated macrophages exhibit phenotypic heterogeneity and immunosuppressive tumor-immune interaction signature is associated with poor survival

ccRCC is recognized as highly immune infiltrated tumor with a dynamic microenvironment. The compositional changes that occur along tumor stage progression^21^ and in response to immunotherapy treatment^22,35^ have a profound impact on patient survival. Therefore, the phenotypic states of immune populations represent potentially druggable targets for advanced and metastatic ccRCC treatments.

Within the immune compartment, we identified all major lymphoid and myeloid cell populations including plasma cells (*IGKC*, *IGHG1*), B cells (*CD79A*, *MS4A1*), mast cells (*TPSB2*), NK cells (*GZMB*, *NKG7*), classical (*CD14*) and non-classical (*FCGR3A*) monocytes and two major groups of T cells and macrophages (Figure 1b), in concordance with previous ccRCC studies^18,19,21^. As expected, the tumor samples were enriched in tumor-associated macrophages (TAMs) that clustered into four transcriptionally distinct subpopulations (Figure 2a). The TAM 1 and TAM 2 cells expressed genes hinting towards M1 and M2 polarization, respectively (Figure 2a), thus encompassing a traditional view of TAM dichotomy. However, TAM 3 and TAM 4 subpopulations did not follow a clear activation pattern, despite their marker genes seemed to reflect an alternatively activated macrophage phenotype (Figure 1f, Supplementary file 1). For example, while the expression of certain immunosuppressive genes, such as *MARCO*, were clearly diminished in TAM 3/4 cells, other immune-response modulating genes such as *VSIG4*^36^ or *VSIR* were highly expressed in TAM 4 population. In addition, among all TAM populations, TAM 4 demonstrated the highest expression of complement system C1Q genes (Figure 2a), products of which are known to promote tumor progression in ccRCC by interacting with tumor-produced complement system molecules^37^. Interestingly, some complement components were not only specific to the tumor cells but also present in the stromal compartment, suggesting potential stromal cell involvement in tumor progression (Supplementary figure S2a). These findings support the notion that ccRCC TME is enriched in suppressive macrophages that adapt to the microenvironment-derived signals influencing disease progression^6,10,21^.

The lymphoid compartment predominantly consisted of CD8 T cells (*CD8B*, *DUSP4*), CD4 regulatory T cells (*FOXP3*, *TNFRSF4*), resting/memory T cells (*IL7R*, *CD52*), cytotoxic T cells (*XCL1*, *KLRB1*) and natural killer cells (*GZMB*, *NKG7*). These subpopulations expressed multiple exhaustion markers (Figure 2b), with classic immune-checkpoint molecule *PDCD1* expressed abundantly in CD8 T cell cluster and *CTLA4* enriched in regulatory T cells. The cytotoxic T cell population shared the exhaustion pattern with NK cells characterized by high expression of *CD160*, *EOMES*, *CD38* and *CD69*. As expected, resting/memory T cells displayed the least exhausted phenotype compared to other lymphoid cell populations (Figure 2b). Given the established exhaustion profile of lymphoid cells and immunosuppressive phenotype of myeloid cells^18,21,38^, we evaluated the crosstalk of these immune cell populations and tumor cells.

Receptor-ligand analysis (see Methods) revealed multiple interactions involved in chemokine processing, immune suppression and sustained survival of tumor cells (Figure 2c, Supplementary tables S5, S6). For example, tumor cells were predicted to communicate with monocytes and TAMs through the immune checkpoint *HLA-G – LILRB1/2* axis, which is involved in promoting the immunosuppressive M2 phenotype and immune escape of the tumor^39^. Interestingly, both pro-inflammatory (M1) and anti-inflammatory (M2) TAMs received signals from tumor cells via *SPP1 – PTGER4* interaction, known to promote macrophage polarization towards tumor supporting phenotype in hepatocellular carcinoma^40^. Another important interaction observed in the TME involved T-cell co-stimulatory *CD27 – CD70* axis, targeted at CD8 T cells and CD4 regulatory T cells. Recent studies have shown that this cell-cell interaction is associated with a pro-tumoral effect, primarily driven by chronic stimulation of T cells leading to exhaustion, enhanced survival of regulatory T cells, and recruitment of TAMs^41^. Furthermore, the expression of interaction signature (gene set of both receptors and ligands, Supplementary table S7) was associated with significantly lower overall survival (Figure 2d, Supplementary table S8) and steadily increased along the progression of the disease in the TCGA KIRC dataset (Figure 2e). Therefore, our analysis of the ccRCC TME reveals the extensive network of immune and cancer cell interactions that are involved in establishing an immune-suppressive TME for sustained tumor survival and growth.

### Tumor endothelial cells are diverse and play a role in re-shaping the tumor microenvironment, associated with worse overall survival

The highly vascularized appearance of ccRCC tumors is often attributed to the abnormal accumulation of hypoxia-inducible factors^2,3^ that create pseudohypoxic conditions and subsequently increase production of angiogenic factors. To this day, the heterogeneity and possible regulatory role of the tumor vasculature in ccRCC remains poorly described. Focusing on ccRCC endothelium in our scRNA-Seq dataset we identified five tumor vasculature (TV) subpopulations (Figure 3a, c) that were markedly distinct from healthy kidney endothelium (Figure 3b) and featured upregulation of genes important in vascularization, angiogenesis and disease progression. For instance, among the multiple overexpressed genes (Supplementary table S9), the TV cells displayed elevated levels of the fenestration marker *PLVAP,* which is recognized as a therapeutic target in hepatocellular carcinoma^42^; *ANGPT2,* which stimulates angiogenesis in autocrine manner and is involved in recruitment of immunosuppressive TAMS^43^; *IGFBP7,* which is clinically used acute kidney injury urinary biomarker^44^. Moreover, endothelial migration stimulating insulin receptor (*INSR*) was overexpressed in tumor endothelium and is known to be associated with poor overall survival in bladder cancer, which, similarly to ccRCC, is frequently resistant to VEGF pathway targeted therapy^45^. These findings highlight the abnormal, fenestrated nature of tumor endothelial cells and might provide future guidance for tumor-specific vasculature identification in ccRCC.

**Figure 3.**
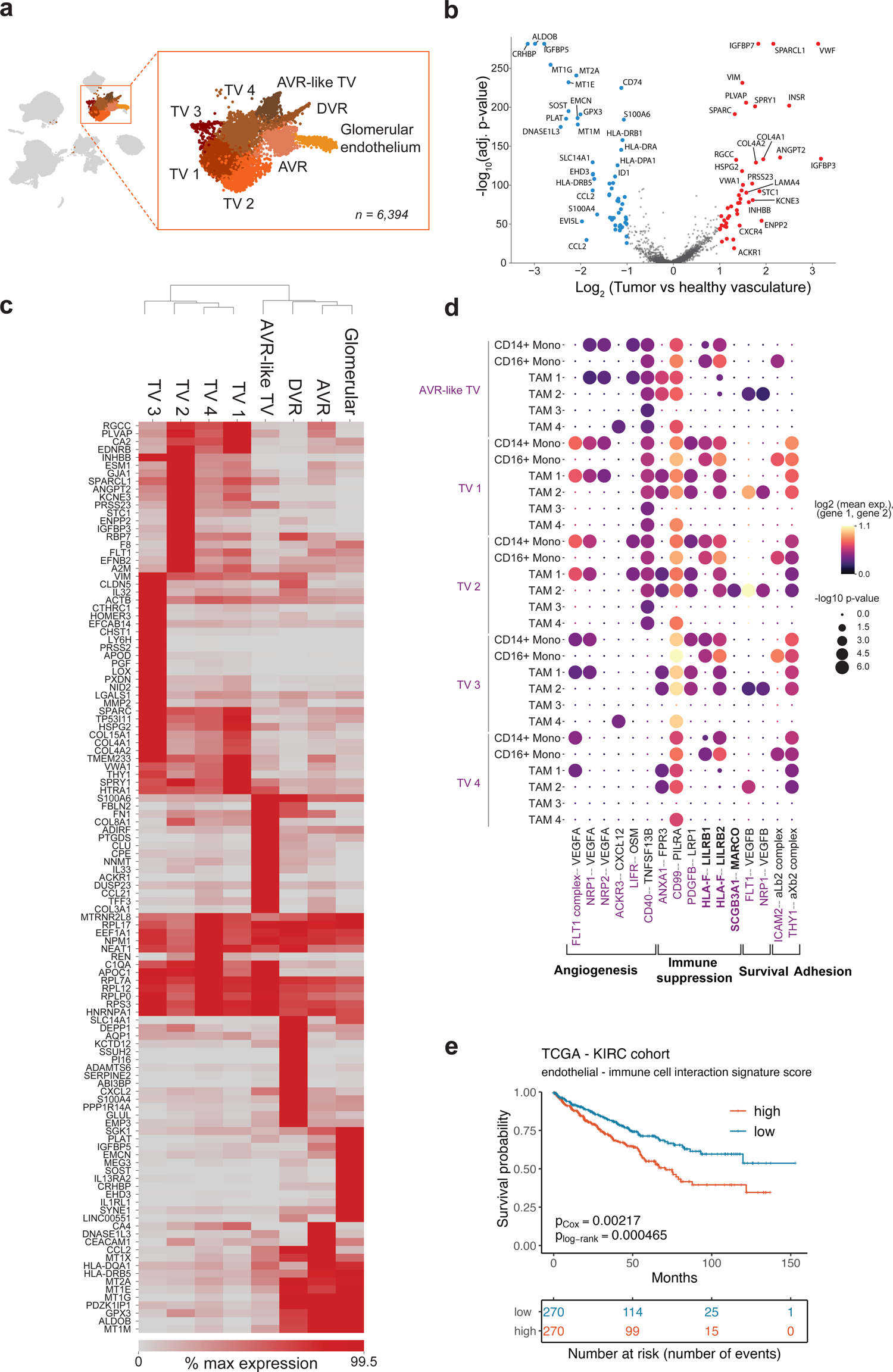
Assessing the heterogeneity of tumor vasculature of ccRCC. a) A close-up of endothelial cell subpopulations. b) Tumor and healthy vasculature comparison shows upregulation of angiogenesis related genes in tumor vasculature. c) Differential gene expression between vasculature subpopulations. Only genes with Benjamini-Hochberg adjusted p-value <0.05 are shown. d) Tumor endothelium and myeloid cells demonstrate abundant cell-cell interactions. e) Collective tumor vasculature – immune cell communication signature expression is associated with a worse overall survival in TCGA KIRC dataset. AVR – ascending vasa recta, DVR – descending vasa recta, TV – tumor vasculature.

Within the tumor vasculature we found an ascending vasa recta-like population that was transcriptionally closer to the healthy endothelium cells than to other tumor vasculature cells (Figure 3c), as noted in previous work^15^. Intriguingly, our ccRCC atlas also unveiled a novel, uncharacterized population of tumor vasculature (referred to as TV 3) that appeared as the most distinct from the rest of TV cells (Figure 3c). This population was marked by high expression of tip cell markers *LOX*, *PXDN*, *LY6H* and *PGF*^31,46^ (Supplementary figure S3, Supplementary table S10), indicative of an invasive phenotype. Furthermore, TV 3, along with TV 1 and TV 4, displayed elevated expression of extracellular matrix constituents, including pro-angiogenic and potentially pro-metastatic collagen type IV and perlecan (*HSPG2*) (Figure 3c)^47–49^. Meanwhile, TV 2 overexpressed multiple genes implicated in tumor progression, such as VEGF receptor *FLT1*, *ESM1*, *ANGPT2*, *KCNE3*, coagulation factor VIII (*F8*) (Figure 3c), which are involved in tumor-associated angiogenesis^49,50^. In addition, TV 2 was marked by high expression of autotaxin (*ENPP2*), a potent stimulator of tumor development and invasion, which has been associated with acquiring resistance to the antiangiogenic drug sunitinib in ccRCC^51^ (Figure 3c). Interestingly, a fraction of cells from all tumor vasculature sub-populations expressed *INHBB* and *SCGB3A1* (Supplementary figure S3), which, in concert with perivascular *TNC* (in our dataset expressed by myofibroblasts, Figure 5b), have recently been demonstrated to orchestrate the pro-metastatic niche in lung metastasis models in mice^52^. Thus, the tumor vasculature in ccRCC appears to be highly heterogeneous and expresses a variety of angiogenesis-related and tumor-promoting factors.

Subsequently, we investigated the potential interactions between tumor vasculature and other cell types within the TME. Cell-cell communication analysis using CellPhoneDB^53^ revealed crosstalk between vascular and immune cells involved in angiogenesis, immune suppression and adhesion (Figure 3d, Supplementary figure S2b). Unexpectedly, our analysis revealed that tumor vasculature delivers immunosuppressive signals previously thought to be confined to the tumor cells, such as the interactions between *TIGIT* and *NECTIN2* (Supplementary figure S2b) or *HLA-F* and *LILRB1/2* (Figure 3d). Also, we observed several known interactions mediated by myeloid cell produced TNF-α with tumor endothelium i.e *TNF*

*– NOTCH1* (Supplementary figure S2b), which induces *JAG1* expression and enhances migration and proliferation of endothelial cells upon subsequent VEGF exposure^54^. Importantly, a higher degree of cell-cell communication between tumor vasculature and immune cells, as evaluated by higher expression of receptor and ligand pairs, was found to result in a significantly lower overall survival in TCGA KIRC cohort (Figure 3e).

These findings suggest notable tumor vasculature participation in tumor progression and tumor microenvironment shaping through the expression of angiogenesis-related genes, tumor-promoting extracellular matrix molecules, and active immunosuppressive communication with immune cells.

### A novel subpopulation of tumor endothelium expresses genes involved in epithelial-mesenchymal transition associated with worse patient survival

The novel tip cell-like tumor vasculature population (TV 3 in Figure 3a) expressed *LOX*, *PXDN*, *LY6H* and *PGF*, which are not only denoted as tip cell markers, but have also been implicated in tumor growth promotion within the TME. For example, placental growth factor (*PGF*), a member of VEGF family, can directly interact with VEGF receptors and increase vascular permeability while promoting M2 macrophage polarization^55^. In *PGF*-deficient mice, tumor-associated M1 macrophage polarization is largely restored while tumor vasculature appears normalized^56^. Lysil oxidase *LOX* and peroxidase *PXDN* are involved in cross-linking of the collagen type IV rich extracellular matrix and basement membrane, which is essential for growth factor induced endothelial cell proliferation and survival^57^. Inhibition of ECM cross-linking through lysil oxidase knockdown has been shown to impair vessel sprouting^31^. Therefore, the tumor vasculature 3 population represents the leading tip cell phenotype in angiogenic sprouting and is potentially involved in promoting tumor progression.

Molecular Signatures Database Hallmark gene set over-representation analysis in tumor, tumor vasculature and stromal cell populations (top 100 marker genes) revealed, as expected, hypoxia and glycolysis terms in tumor cells (Figure 4a, Supplementary table S11). However, this analysis also uncovered an enrichment of epithelial-mesenchymal transition (EMT) associated genes in all tumor vasculature and stromal cell subpopulations. Interestingly, the overexpression of EMT pathway overlapping genes for AVR-like tumor vasculature (Figure 4b) and TV 3 population (Figure 4c) was associated with a significantly worse overall survival in the TCGA KIRC cohort. In this context, it is important to note that the specific genes overlapping with the EMT differed between these subpopulations (Supplementary table S12). Also, even though other cell populations, such as stromal cells and the rest of tumor vasculature had a significant overlap with the EMT pathway (Supplementary figure S4a), no effect on patient survival in the TCGA KIRC cohort was observed (Supplementary figures S4b-g). Overall, our findings highlight the presence of a tip cell-like tumor endothelium subpopulation associated with an aggressive phenotype, potentially influencing ccRCC disease progression and survival.

**Figure 4.**
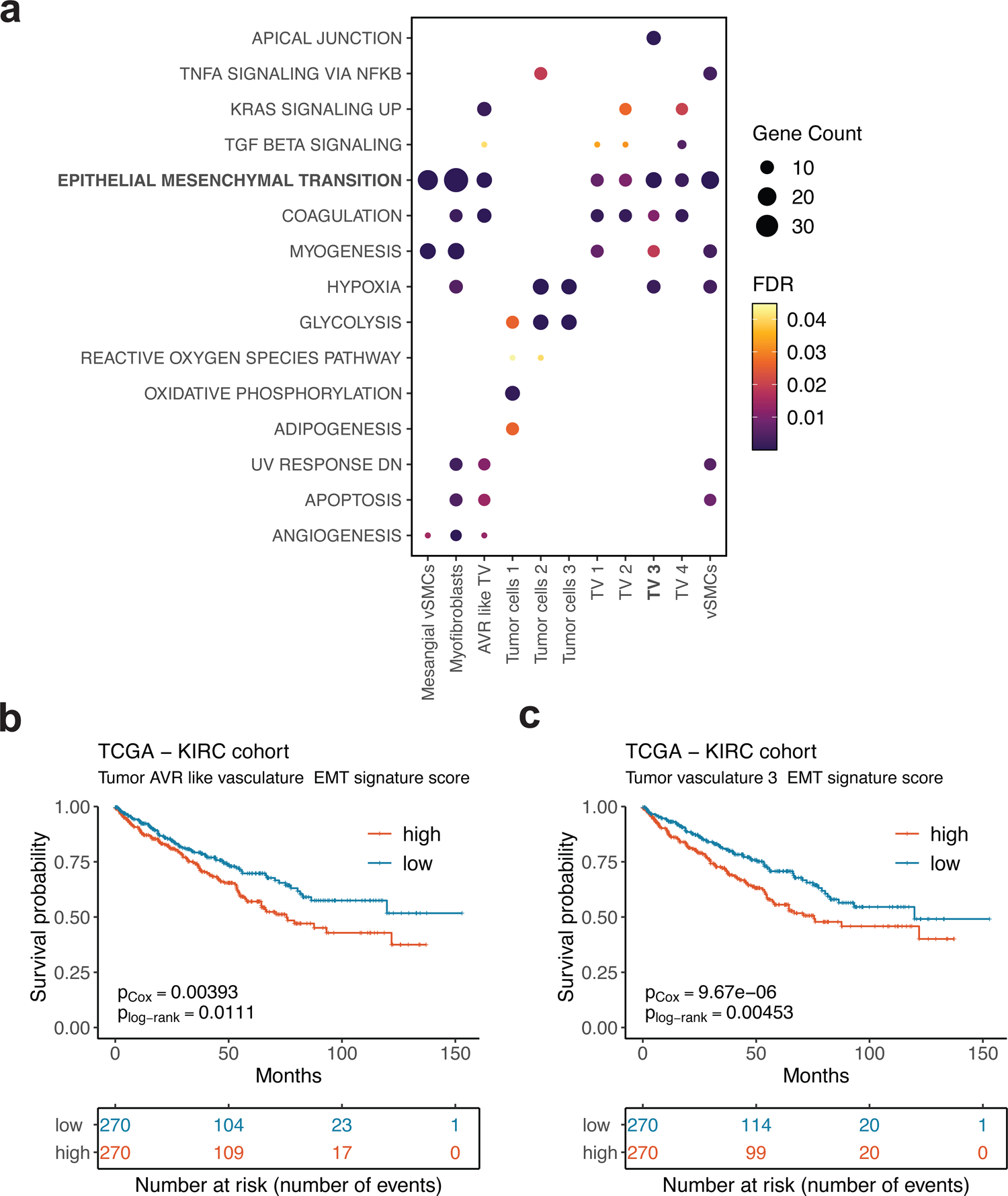
MSigDB Hallmark pathway overrepresentation analysis. a) Tumor vasculature and stromal cell populations are enriched in epithelial-mesenchymal transition (EMT) signature. b) Tumor AVR-like vasculature and c) tip-like tumor vasculature 3 signature genes overlapping with EMT pathway associate with worse overall survival in the TCGA KIRC cohort.

### Stromal cells remodel the ECM and potentially contribute to immunosuppression of TAM populations

Finally, we investigated the putative roles of stromal cells in the ccRCC tumor microenvironment. While stromal cells have been recognized as important components of the TME^34^, their specific contribution in ccRCC have received much less attention compared to immune or tumor cells. Graph-based clustering of our dataset revealed three cell populations within the stromal cells: vascular smooth muscle cells (vSMCs), myofibroblasts and mesangial/vSMCs (Figure 5a, b, Supplementary table S13). The vSMCs expressed markers *TAGLN*, *ACTA2* and *MYH11*, while myofibroblasts were enriched for ECM constituents (Collagen types I, III, IV, VI and fibronectin) including markers *TIMP1* and *ACTA2* (Figure 5b). The precise annotation of the third stromal cell population was challenging due to simultaneous upregulation of mesangial marker *PDGFRB* and vSMC genes (Supplementary file Table 1). Interestingly, this population featured substantial transcriptional differences between tumor and healthy tissue (Supplementary figure S5, Supplementary table S14). In tumor samples, the mesangial/vSMC population overexpressed tumor marker *NDUFA4L2* as well as some stress-related genes, such as *CD36*, which is upregulated in chronic kidney disease and associated with poor prognosis in ccRCC^58,59^, and renin (*REN*), which is expressed by mesangial cells under disturbed homeostasis^60^ (Supplementary figure S5). Thus, it appears that the mesangial/vSMC population is reactive to the disruptive microenvironmental changes exerted by the tumor.

**Figure 5.**
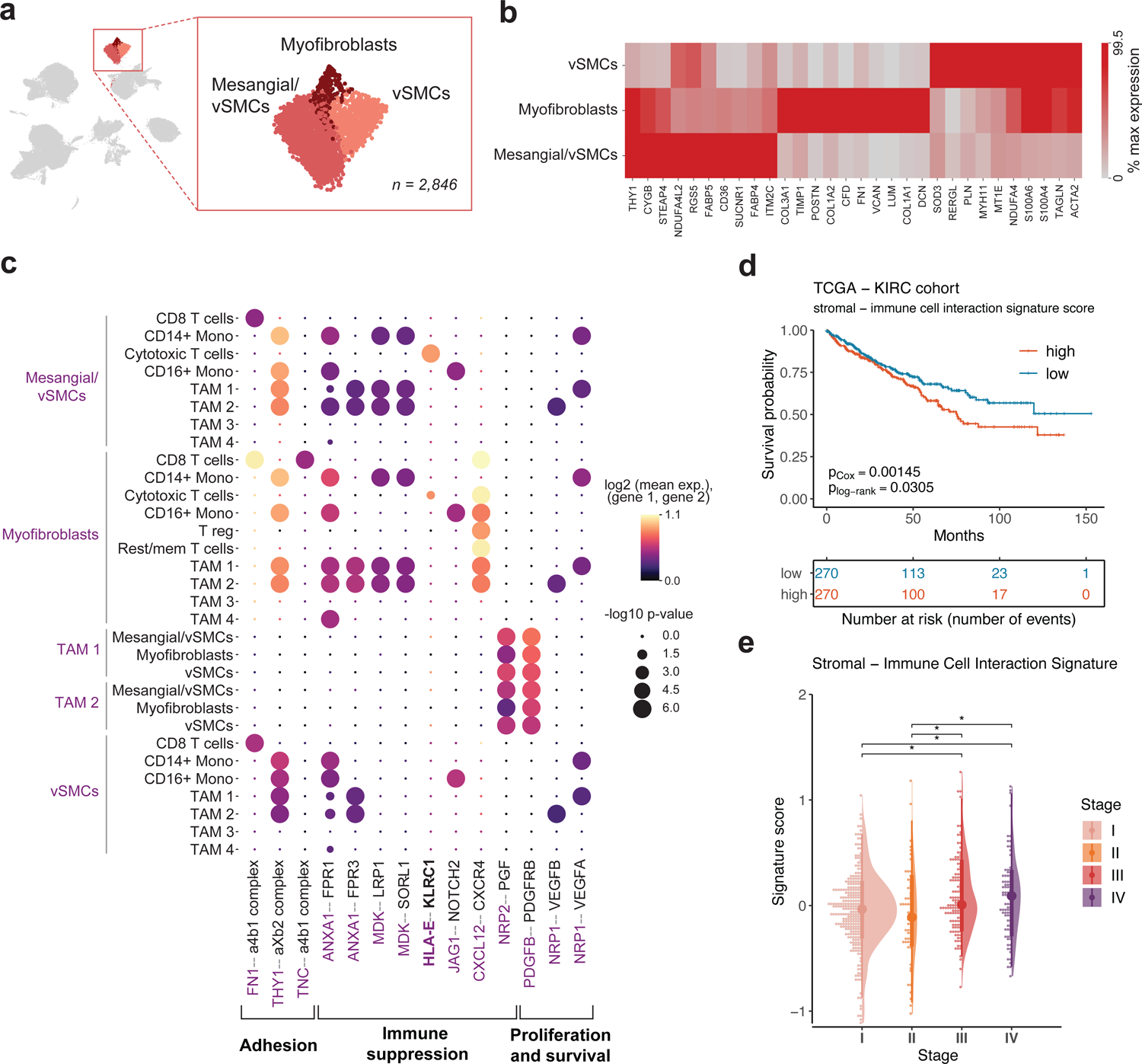
Assessing the heterogeneity of stromal cells in the TME. a) Stromal cell populations consisting of vSMCs, myofibroblasts and mesangial/vSMCs. b) Differential gene expression between stromal cell subpopulations. Only genes with Benjamini-Hochberg adjusted p-value <0.05 are shown. c) Stromal and immune cells exhibit immunosuppressive interactions mediated by stromal cells. d) Expression of collective stromal-immune cell interaction signature gene set associates with worse overall survival in the TCGA KIRC cohort. e) Stromal-immune cell interaction signature expression increases along the progression of the ccRCC disease. vSMCs – vascular smooth muscle cells.

Cell-cell interaction analysis between stromal and immune cells revealed putative interactions related to stromal cell proliferation and survival, as well as immune cell suppression and adhesion. Majority of immunosuppressive signals originating from the stromal cells were directed at TAM 1 and TAM 2 subpopulations (Figure 5c). For instance, we identified *ANXA1 – FPR1* interaction, which is involved in anti-inflammatory macrophage polarization and tumor progression in various cancers^61,62^. Furthermore, we found an indication of myofibroblast and mesangial/vSMC communication with cytotoxic T cells via *HLA-E – KLRC1*, which has recently been proposed as a new targetable path of T cell exhaustion in bladder cancer^63^. Treatment of *HLA-E* positive tumors with anti-KLRC1 antibodies has shown a strong effect in restoring the anti-tumor immunity^64^. Interestingly, our analysis shows that this communication signature is associated with worse overall survival in the TCGA KIRC dataset (Figure 5d), and the expression of genes involved in the stromal-immune cell communication increased with advancing stage of the disease (Figure 5 e). Collectively, our results suggest that stromal cells are actively involved in modulating the tumor microenvironment in ccRCC through therapeutically relevant paths.

## DISCUSSION

The single-cell transcriptomic studies have provided valuable insights about the origin of ccRCC^14,15^, malignancy programs of the tumor^16^, immune cell population phenotypical changes during tumorigenesis^21^ and immunotherapy treatment^18,22^ among other. Complementing these ongoing efforts to better characterize ccRCC tumor microenvironment we profiled single-cell transcriptomes of human ccRCC tumor samples along with healthy adjacent tissues. In contrast to previous studies that used cell enrichment prior to scRNA-Seq, our strategy relied on a rapid isolation of cells from ccRCC specimens, without involving any type of sorting or cell enrichment. As a result, we could capture a rich diversity of cells constituting heterogeneous TME that were either significantly depleted or absent in previous studies. Given that immune compartment in our dataset largely recapitulated previous findings^17–22^, we mainly focused on the phenotypic heterogeneity and cellular interactions of the often overlooked and underappreciated endothelial and stromal cell populations.

Endothelial cells are very important in ccRCC tumorigenesis and to this day remain the main targets of therapeutics in advanced and metastatic disease^2^. The tumor endothelial cells identified in our study include a novel, previously uncharacterized tip cell phenotype, enriched for epithelial-mesenchymal transition pathway genes that are associated with poor overall survival. Indeed, the previous single-cell studies in ccRCC have also captured endothelial cells, however, these were most often represented by two major phenotypic subpopulations that are also found in our ccRCC atlas. For instance, *Zhang et al.*, reported *ACKR1*+ and *EDNRB*+ endothelium, while *Long et al.* reported *VCAM1*+ and *VCAM1*-vasculature populations. Consistently, in our dataset we find a population co-expressing ascending vasa recta marker *ACKR1* and *VCAM1* (tumor AVR-like vasculature), however, *EDNRB* is expressed by tumor vasculature 1, 2, and 4 populations, but not by tumor vasculature 3 (Supplementary figure S3), further supporting that this endothelial (*PECAM1*+) phenotype has not been characterized in ccRCC.

The tip cell population (TV 3) in our dataset shares similarities with a tip cell population observed in lung cancer (*LOX*, *PXDN*, *PGF*, *LXN*, collagen type IV enriched) where it was shown to correlate with worse patient survival^31^. The authors have found this phenotype the most congruent across several species and tumor types, including kidney cancer (as determined by bulk proteomics), which raises a question about why previous single-cell studies of ccRCC did not capture this rare population. Furthermore, the authors demonstrated that tip cell marker *LOX* knock-down impaired vessel sprouting, suggesting that the reported population in ccRCC might be of interest for future research as a potential therapeutic target.

In line with our findings, *Long et al.,* showed that *VCAM1*+ population (labeled as AVR-like tumor vasculature in our dataset) is enriched for EMT signature^16^, yet our pathway over-representation analysis indicates similar association with EMT for all tumor vasculature and stromal cell populations, not just the AVR-like population (Figure 4a). On another hand, the worse overall survival in association with EMT was pronounced only for AVR-like and the tumor vasculature 3 populations, further emphasizing the diversity of tumor endothelial cells and potential importance of the reported tip cell phenotype. *Alchahin et al.,* also reported association with EMT for endothelial and stromal cells, but did not discriminate healthy kidney and tumor endothelial cells. On the contrary to our findings, they report lower endothelial cell abundance in tumor samples as compared to healthy tissues^20^. Such discrepancies between different studies can be related to technical aspects, for instance, processing of the samples, and further underline the importance for accurate phenotypic characterization of the tumor vasculature cells in ccRCC.

Our findings suggest two major modes of action of the tumor vasculature cells in the TME. First, remodeling of the ECM by active deposition of various ECM constituents and expression of their modifying agents related to EMT (i.e. *LOX*, *PXDN* in tumor vasculature 3) and second, active engagement in cellular communication in the tumor microenvironment, mostly involved in immune suppression and angiogenesis maintenance. Interestingly, spatial transcriptomic profiling of ccRCC by *Li et al.,* showed that collagen producing endothelial cells localize at the tumor-normal interface enriched in EMT-high tumor cells and *IL1B*+ macrophages^17^. These findings are also corroborated by our results suggesting that tumor endothelial cells might indeed contribute to EMT in ccRCC and interact with TAMs. The cell-cell communication analysis uncovered diverse interactions of clinical relevance enriched in the tumor vasculature and stromal cell communication with immune cells (Figure 3d, 5c). For instance, in 2021, a phase I-II clinical trial (ID NCT04913337) began for LILRB1 and LILRB2 inhibitor as a monotherapy or in combination with Pembrolizumab (anti PD-1) for advanced or metastatic solid tumors, including ccRCC. Inhibition of LILRB2 reprograms myeloid cells to a stimulatory (pro-inflammatory) state, while inhibition of LILRB1 stimulates the reprogramming of both myeloid and lymphoid cells. Our analysis suggests that *LILRB1/2*+ immune cells interact not only with tumor cells, but also with endothelial cells. Similarly, endothelial cell-expressed *NECTIN2* associated with *TIGIT* expressed by regulatory T cells, an interaction that has gained increased attention over the last few years and is currently exploited in a multitude of clinical trials^65^. Another intriguing interaction observed between TV 2 and TAM 2 populations was *SCGB3A1 – MARCO*. As demonstrated recently, *SCGB3A1*, a secreted secretoglobin family member produced by endothelial cells, is a crucial component of a pro-metastatic niche and induces stem cell properties in cancer cells, while macrophages are also required for the niche maintenance^52^. However, *SCGB3A1 – MARCO* interaction in ccRCC, to our knowledge, has not been described.

It is worth emphasizing that stromal cells in our dataset were involved in communication with immune cells in a suppressive manner, suggesting their participation in maintaining a pro-tumorigenic niche, especially considering the difference of mesangial/vSMCs population expression in tumor vs healthy adjacent tissue. Moreover, the communication signature expression associated with worse overall survival and increased along the progression of the disease in the TCGA KIRC dataset. On a side note, increase of stromal cells has recently been shown in recurrent RCC as compared to primary disease, furthermore, stromal cell-produced Galectin-1 (*LGALS1*) inhibitor significantly reduced tumor mass and improved anti-PD-1 immunotherapy efficacy in murine models^66^. Another report showed that co-targeting stromal cells expressing PDGFRs and endothelial cells expressing VEGFRs delays tumor vascularization and has clinical efficacy in pancreatic neuroendocrine tumors^43^. Therefore, there is a need for in-depth characterization of ccRCC stromal cells and further validation of their pro-tumorigenic properties. Understanding the role of stromal cells in the TME could provide valuable insights for the development of targeted therapies.

Overall, our study introduces an invasive tumor-associated endothelial tip cell phenotype and provides new insights into the characterization of the TME in ccRCC. We propose that tumor endothelial cells favor tumor progression and potentially metastatic dissemination through the expression of metastasis promoting factors, specific extracellular matrix components and indirectly via targetable interactions with immune cells in the TME. Undoubtedly, future functional studies are needed to elucidate the exact roles of the described diverse tumor endothelial cells and explore their potential as therapeutic targets in ccRCC.

## MATERIALS AND METHODS

### Sample acquisition

Fresh ccRCC tumor (n=8) and healthy-adjacent (n=9) paired kidney tissues were obtained from the National Cancer Institute (Vilnius, Lithuania) with a bioethics committee approval No.2019/2-1074-586. No patient had received prior systemic therapy for their cancer. Samples were collected during an open or laparoscopic, partial or radical nephrectomy surgery, placed on ice and rapidly (<1 hour) transferred to the laboratory for dissociation. Sample T1 (tumor from patient P1) was highly necrotic, thus excluded from analysis. Clinical characteristics of all samples profiled are provided in Supplementary Table S1.

### Sample processing

Sample preparation was performed according to the scRNA-Seq protocol^67^, yet without FACS-based enrichment. Briefly, patient derived tumor tissues were dissociated using Tumor Dissociation Kit (Miltenyi Biotec, cat.no.130-095-929) in an automated instrument gentleMACS Octo Dissociator with Heaters (Miltenyi Biotec) as per manufacturer’s instructions. Healthy-adjacent tissues were dissociated using Tissue Dissociation Kit I (Miltenyi Biotec, cat.no. 130-110-201). After dissociation, red blood cells were removed from the samples using RBC lysis reagent (Miltenyi Biotec, cat.no.130-094-183). After RBC lysis, cells were washed 3 times in ice-cold 1X DPBS (Gibco, cat.no. 14080-048) at 500g for 5 min. Cell viability and count was assessed using Trypan Blue dye (Gibco, cat.no. 15250061) on a hemocytometer. No further enrichment or selection of cells was performed. Cell suspension was immediately loaded onto inDrops platform^68^ for cell barcoding experiment.

### Single cell barcoding, library preparation and sequencing

Dissociated cells were isolated in 1 nanoliter droplets and their transcriptomes barcoded using a modified version of inDrops protocol^69^. Specifically, instead of linear cDNA amplification by *in vitro* transcription we used template switching and PCR amplification. For that purpose, we isolated the cells at occupancy 0.1 alongside with barcoding beads (Atrandi Biosciences, cat.no. DG-BHB-C) and reverse transcription/lysis mix, the latter supplemented with a template switching oligonucleotide, TSO (see Table 2 for composition). We used cell barcoding chip (Atrandi Biosciences, cat.no. MCN-05) to inject the cells, DNA barcoding beads, and RT/lysis mix at flow rates of 250, 60, 250 µl/hr, respectively. The droplet stabilization oil (Atrandi Biosciences, cat. no. MON-DSO2) was set at 700 µl/hr. The emulsion was collected off-chip on ice rack and briefly exposed to UV light (5 min at 6.5 J/cm^2^ of 350 nm, Atrandi Biosciences, cat.no. MHT-LAS2) to release the photo-cleavable RT primers from the barcoding hydrogel beads. The RT reaction was performed at 42 °C for 60 min followed by 5 min at 85 °C. The post-RT emulsion was burst with 10% emulsion breaker (Atrandi Biosciences, cat.no. MON-EB1) and pooled material was used for subsequent library construction.

**Table 2.** Lysis/RT reaction mix for single-cell mRNA barcoding.

| Reagent | Amount, μl | Concentration in droplet |
| --- | --- | --- |
| Nuclease-free water | 21 | --- |
| 5X RT buffer | 60 | 1X |
| TSO primer (0.5 mM) | 15 | 25 µM |
| dNTP (10 mM each) | 15 | 0.5 mM |
| 10% (v/v) NP-40 (lysis agent) | 9 | 0.3 % |
| RiboLock RNase Inhibitor | 15 | 1 U/ul |
| Maxima H minus RT enzyme | 15 | 10 U/ul |
| Total volume | 150 | --- |

### Library construction

The barcoded-cDNA was purified twice with 0.8X AMPure XP reagent (BeckMan Coulter, cat.co. A63881) as per manufacturer’s instructions. Next, cDNA was PCR amplified with KAPA HiFi Hot Start Ready Mix (Roche, cat.no. KK2601) using cDNA FWD primer and cDNA REV primers (see Table 3). Amplified DNA was fragmented and ligated to adapter using instruction and reagents provided by NEBNext® Ultra™ II FS DNA Library Prep (NEB, cat.no. E7805S). Finally, the libraries were amplified by 12-rounds of indexing PCR (2X KAPA HiFi Hot Start Ready Mix, Roche, cat.no. KK2601). Library quality was assessed using Bioanalyzer DNA High Sensitivity chip (Agilent, cat.no. 50674626). The libraries were sequenced on Illumina NextSeq 550 platform in multiple batches using either NextSeq 500/550 High Output Kit v2.5 (75 Cycles) (Illumina, cat.no. 20024906) or NextSeq 500/550 High Output Kit v2.5 (150 Cycles) (Illumina, cat.no. 20024907).

**Table 3.**
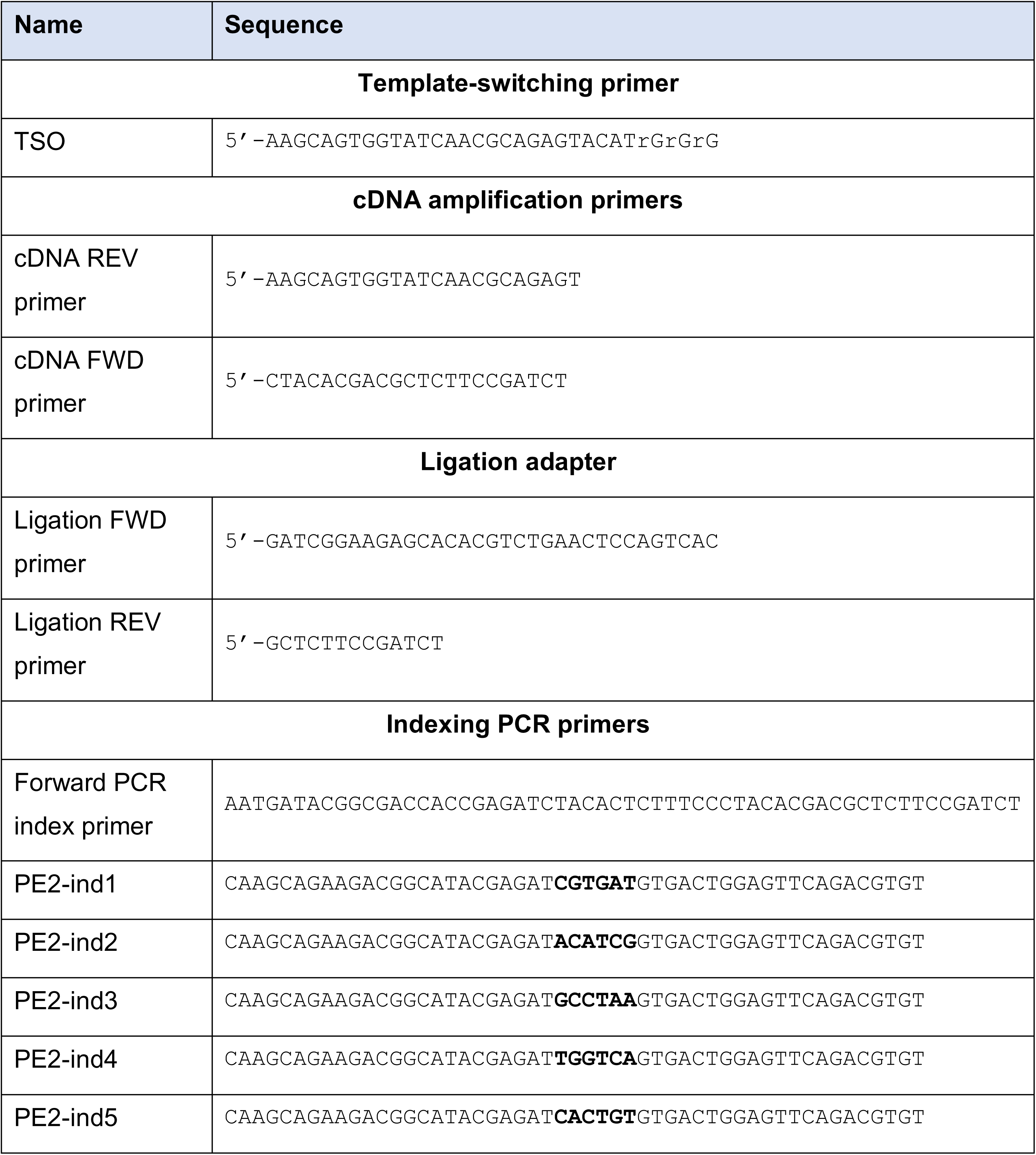
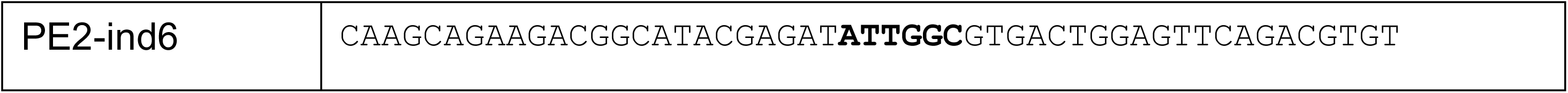
List of DNA oligonucleotides.

### Raw sequencing data processing

The STARsolo pipeline (https://github.com/jsimonas/solo-in-drops) was used to process the data and to obtain expression matrices. STAR (version 2.7.6a) was run with the following parameters: --soloMultiMappers Uniform, --soloType CB_UMI_Simple, --soloUMIfiltering MultiGeneUMI, and --soloCBmatchWLtype 1MM. Homo sapiens (human) genome assembly GRCh38 (hg38) and Ensembl v93 annotations were used as the reference.

### Data analysis: quality control, doublet and RBC removal

Starting with cell x gene matrices, analysis was performed in Python using scanpy toolkit (Table 4). All notebooks are provided at https://github.com/zvirblyte/2023_ccRCC. Briefly, the raw count matrices were uploaded into an AnnData object and filtered by total transcript count and mitochondrial count fraction. The threshold for mitochondrial counts for all libraries was 20%. The total transcript count threshold was determined by evaluating the total count distribution and was selected permissive at minimum 400 UMIs per cell (300 UMIs for libraries T3.1, T9.1, N3.3, N4.3, N2.3). Doublets were removed using Scrublet^70^ (v0.2.3) in the same PCA space used for initial UMAP construction. Scrublet was applied on each emulsion separately. Briefly, the procedure for doublet removal consisted of 1) Calculating doublet scores for each cell in each emulsion using Scrublet; 2) high resolution graph-based clustering using Scanpy’s Louvain algorithm implementation (resolution = 60); 3) evaluation of mean doublet score and fraction of predicted doublets per cluster; 4) manual inspection of doublet-rich clusters in the interactive SPRING application^71^, 5) removal of clusters with high mean doublet score and doublet fraction and no cluster-specific gene expression. This procedure, starting from UMAP construction at step 2) was repeated a total of 2 times and 913 cells (<2% of the total cell population) were removed. Transcriptomes with >1% of total raw counts originating from hemoglobin genes (HBB, HBA1, HBA2, HBD) were considered as red blood cells (RBCs) and 47 such transcriptomes were removed from further analysis.

**Table 4.** Software and algorithms.

| Software | Version | Reference |
| --- | --- | --- |
| solo-in-drops | v1.0 | https://github.com/jsimonas/solo-in-drops |
| STAR | 2.7.6a | https://github.com/alexdobin/STAR,<br>https://doi.org/10.1101/2021.05.05.442755 |
| scanpy | v1.8.0 | <sup>77</sup>, https://scanpy.readthedocs.io/en/stable |
| harmonypy | v0.0.5 | <sup>72</sup>, https://github.com/slowkow/harmonypy |
| scrublet | v0.2.3 | <sup>70</sup>, https://github.com/swolock/scrublet |
| SPRING viewer | N/A | <sup>71</sup>, https://github.com/AllonKleinLab/SPRING_dev |
| scikit-learn | v1.0.2 | https://scikit-learn.org/stable |
| statsmodels | v0.12.2 | https://www.statsmodels.org/v0.12.2 |
| scipy | v1.6.2 | <sup>78</sup>, https://scipy.org |
| anndata | v0.7.6 | https://doi.org/10.1101/2021.12.16.473007,<br>https://anndata.readthedocs.io/en/latest |
| numpy | v1.20.1 | https://numpy.org/doc/1.20/index.html |
| pandas | v1.2.4 | https://pandas.pydata.org |
| louvain | v0.7.1 | https://github.com/vtraag/louvain-igraph |
| umap | v0.5.1 | https://umap-learn.readthedocs.io/en/latest |
| matplotlib | v3.2.2 | https://matplotlib.org/stable/index.html |
| seaborn | v0.11.0 | https://seaborn.pydata.org |
| jupyterlab | v2.2.6 | https://jupyter.org |
| CellPhoneDB | v2.0 | <sup>53</sup>, https://cellphonedb.readthedocs.io/en/latest/index.html |
| R | v4.2.1 | https://www.r-project.org/ |
| tidyverse | v1.3.2 | https://www.tidyverse.org/ |
| biomaRt | v2.52.0 | https://bioconductor.org/packages/biomaRt/ |
| clusterProfiler | v4.4.4 | https://bioconductor.org/packages/clusterProfiler/ |
| TCGAbiolinks | v2.24.3 | https://bioconductor.org/packages/TCGAbiolinks/ |
| survival | v3.3-1 | https://CRAN.R-project.org/package=survival |
| survminer | v0.4.9 | https://cran.r-project.org/package=survminer |

### UMAP construction, clustering and annotation

After filtering and QC steps we retained 50,236 single cells that were used to construct a graph and UMAP representation (Figure 1B). The procedure consisted of 1) normalization to 10 000 total counts, log-transformation and scaling; 2) selection of highly variable genes; 3) PCA; 4) batch correction using Harmony^72^; 5) graph construction and 6) UMAP representation. After normalization, genes with 15 CPTT (counts per ten thousand) in not less than 25 cells were considered abundant and retained, furthermore, mitochondrial and ribosomal genes were excluded and top 2000 abundant and highly variable genes, based on Fano factor (as in ^68^), were used for PCA. To remove batch effects due to different batches of barcoding beads the dataset integration was performed using function scanpy.external.pp.harmony_integrate() with the batch variable ‘beads’. Then, adjacency graph was constructed using sc.pp.neighbors() with n_neighbors=30 and UMAP representation was built using sc.tl.umap() with min_dist=0.4. The resulting representation was used for exploration in interactive SPRING application. Graph-based spectral clustering with varying number of clusters (k) was performed using sklearn.cluster.SpectralClustering() function, the clustering results were explored in the interactive SPRING environment, and k=43 was selected for annotation. Differential gene expression analysis (Mann Whitney U test with Bonferoni-Hochberg correction) was performed and top 25 marker genes for each cluster (adjusted p-value <0.05) were used for in-depth literature analysis and manual cell type annotation (Supplementary file Table 1, Supplementary table S2).

### Sample heterogeneity quantification

To quantify sample heterogeneity, Shannon entropy of samples was calculated for each broad cell category as described in Chan et el.^11^ Briefly, entropy values were calculated for sample frequency in each cell group (stromal, endothelial, tumor, lymphoid, myeloid, epithelial and cycling). To account for differences in the number of cells per group, we subsampled 100 cells from each group 100 times with replacement and calculated the Shannon entropy using function scipy.stats.entropy(). Cells from cluster “Tumor cells 1” were excluded, as they were sample specific.

### Receptor-ligand interaction analysis

Log-normalized expression values for all cell types, excluding healthy epithelial cell populations and cycling cells were used to infer cell-cell interactions using CellphoneDB v.2.0.0^53^ with method “statistical_analysis” and default parameters. Significant (p-value <0.05) cell-cell interactions were explored and selected interactions are shown in Figure 2C, 3D, 5C and Supplementary Figure 2B. Cell-cell interaction signatures for subsequent survival analysis (as in Figure 2D) were constructed by taking both the receptor and ligand genes in the set (provided in Supplementary table S7). Cell-cell interaction analysis results are provided in Supplementary tables S5 and S6.

### Gene set over-representation analysis

Gene set over-representation analysis was employed to evaluate the potential functional significance of a given gene signature. The analysis utilized gene sets obtained from the Hallmark Pathways of the MSigDB database v7.5.1^73^. Gene signatures were then submitted to a hypergeometric test implemented in the enrichGO() function of the clusterProfiler R package^74^ using genes that were detected (nonzero UMI counts) in kidney tissue samples as a universe (background reference). The pathways having FDR (Benjamini-Hochberg) values below 0.05 were considered as significantly over-represented.

### Survival analysis

TCGA KIRC cohort bulk RNA-seq (upper quartile FPKM normalized) and clinical data were downloaded from the NCI GDC Data Portal^75^ using the TCGAbiolinks R package^76^. Cell type signature scoring of the TCGA bulk RNA-seq samples was performed by calculating an arithmetic mean of the z-score transformed expression values for all genes in a given signature. The used gene-wise z-score transformation equalized differences in the gene expression abundances, so that lowly and highly expressed genes would have the same scale and, thus equal weight in the score. The association between signature score and overall survival time was assessed by Kaplan-Meier and multivariate Cox regression analyses. Log-rank tests and Wald tests, respectively, were used to evaluate statistical significance (at level of 0.05) of the performed survival analyses. For the Kaplan-Meier analysis, stratified signature (high - greater or equal than the median signature score; low – lower than the median signature score) was used, while for the multivariate Cox regression analysis, the continuous signature score values were used with patient age and sex as covariates. The survival analyses were conducted using the survival and the survminer R packages.

## DATA AND CODE AVAILABILITY

Upon publication raw data files will be deposited following editorial guidelines. All Jupyter notebooks for scRNA-seq analysis are available at https://github.com/zvirblyte/2023_ccRCC.

## ACKNOWLEDGEMENTS

We are especially grateful to the patients at the National Cancer Institute, Vilnius, Lithuania for participating in this study. This work received funding from European Regional Development Fund [01.2.2-LMT-K-718-04-0002] under grant agreement with the Research Council of Lithuania. The work in S.Ja. group was funded by grant no. S-MIP-17-54. S. Ju was supported by the European Union’s Horizon 2020 research and innovation programme under the Marie Skłodowska-Curie grant agreement no. 101030265. We are grateful to Karolis Goda for wet-lab assistance, Rapolas Zilionis for valuable discussions and input on data analysis, and the members of the Oncourology Department at the National Cancer Institute (Lithuania) for their valuable support and kind assistance.

## AUTHOR CONTRIBUTIONS

J.Z., J.N. single-cell RNA-seq experiments, library preparation and sequencing; J.N., R.K. biospecimen logistics and processing; M.K., A.U., patient consent, biospecimen curation, acquisition and logistics; J.Z. data analysis and interpretation, initial manuscript draft; S.Ju. data management and analysis; J.N., S.Ju., S.Ja., L.M. proofreading; J.Z., L.M. manuscript revision and preparation; S.Ja., and L.M. study design and funding acquisition; L.M. supervision. All authors have read and approved the final manuscript.

