## Supplementary file 1 for "Single-cell transcriptional profiling of clear cell renal cell carcinoma reveals an invasive tumor vasculature phenotype"

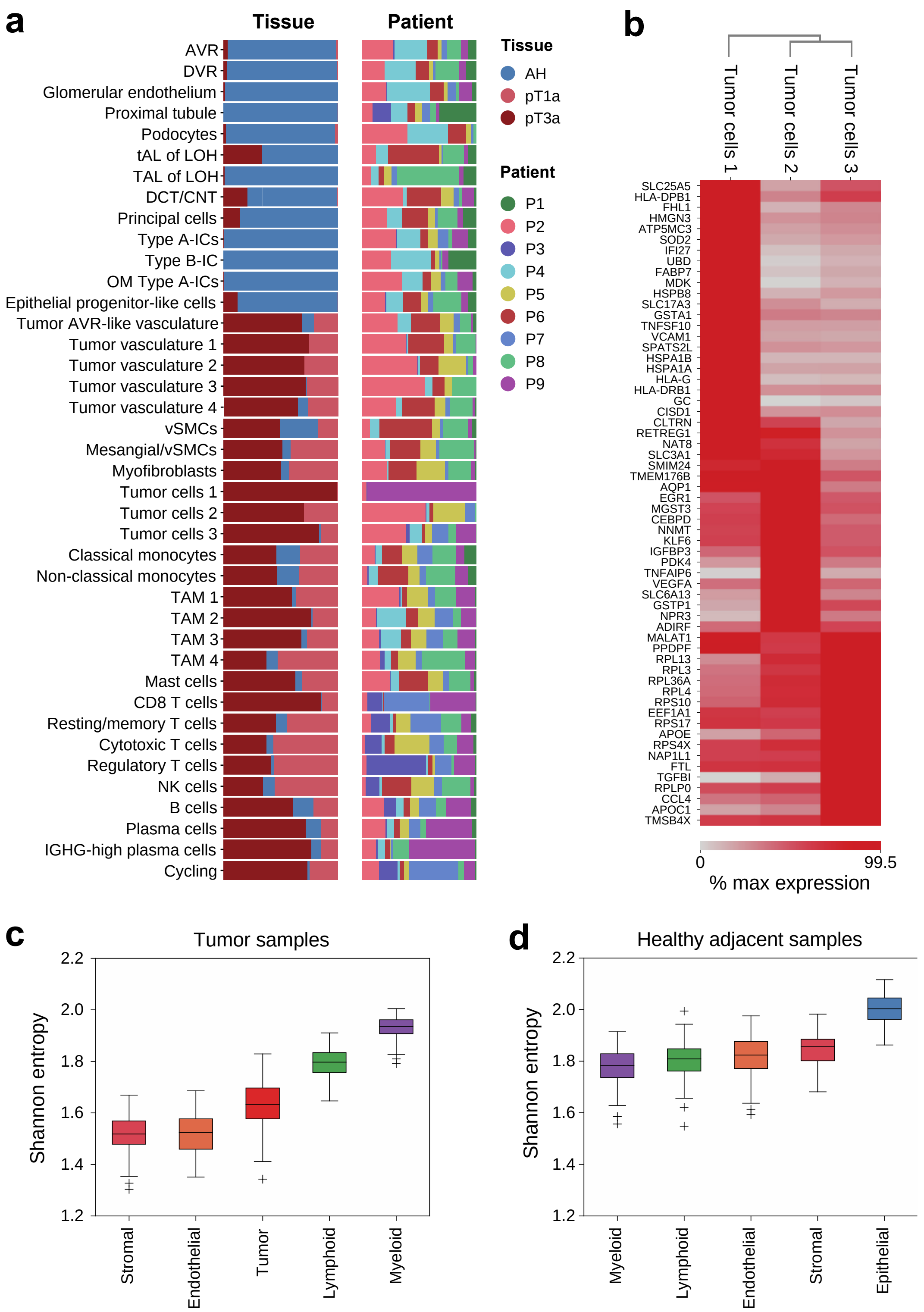


**Supplementary figure S1.** Cell composition and inter-patient variability in ccRCC. **a**) Cell composition by disease stage and patient ID. Specialized epithelial and endothelial cells originate mostly from the healthy-adjacent tissues, while immune, tumor, endothelial and stromal cells are enriched in the tumor samples. Different types of cells are adequately represented by multiple samples, except for tumor cells 1 population, which appears specific to patient P9. **b)** Differential gene expression between stromal cell subpopulations. Only genes with Benjamini-Hochberg adjusted p-value <0.05 are shown. **c, d**) Tumor and healthy adjacent sample heterogeneity for broad cell group as measured by Shannon entropy. Lower entropy values indicate higher sample heterogeneity. AH – adjacent healthy.

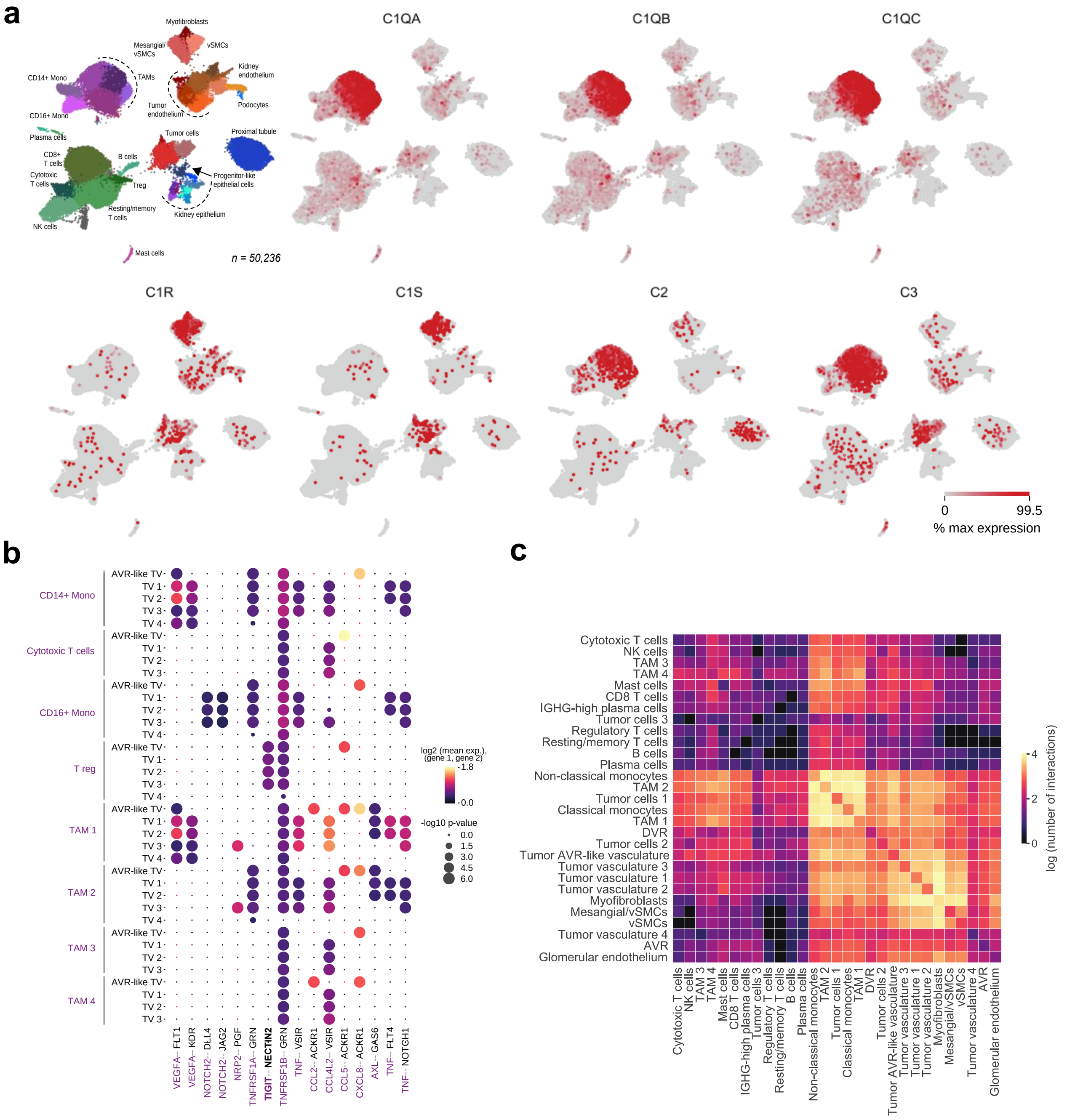

**Supplementary figure S2.** Expression of complement system genes and cell-cell communication analysis. **a)** Expression of complement system molecules C1QA, C1QB and C1QC is specific to tumor associated macrophages, while C1R, C1S are expressed by tumor and stromal cells. **b)** Cell-cell communication analysis between immune cells and tumor vasculature reveal immunosuppressive TIGIT-NECTIN2 interaction between tumor vasculature and regulatory T cells. **c)** Count matrix of all major cell-cell interactions observed within the TME of ccRCC

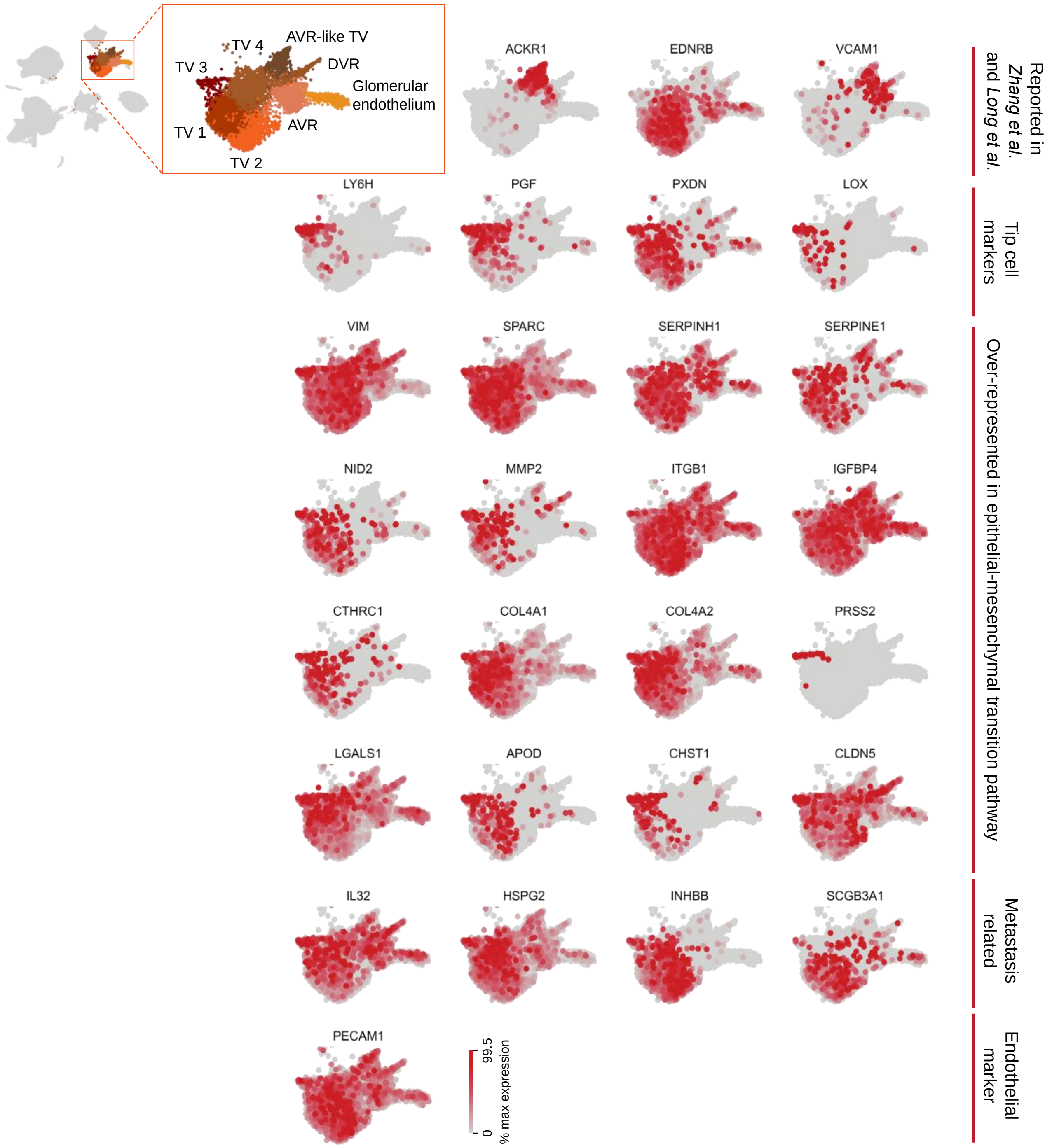


**Supplementary figure S3.** Expression of metastasis associated genes, tip-cell markers and genes overlapping with epithelial-mesenchymal transition pathway in tumor vasculature. The tip-cell markers are mostly enriched in tumor vasculature 3 population, while other genes are expressed in multiple tumor vasculature populations in heterogeneous manner. Previously reported markers *ACKR1*, *EDNRB* and *VCAM1* are not expressed in tumor vasculature 3. AVR – ascending vasa recta, DVR – descending vasa recta, TV – tumor vasculature.

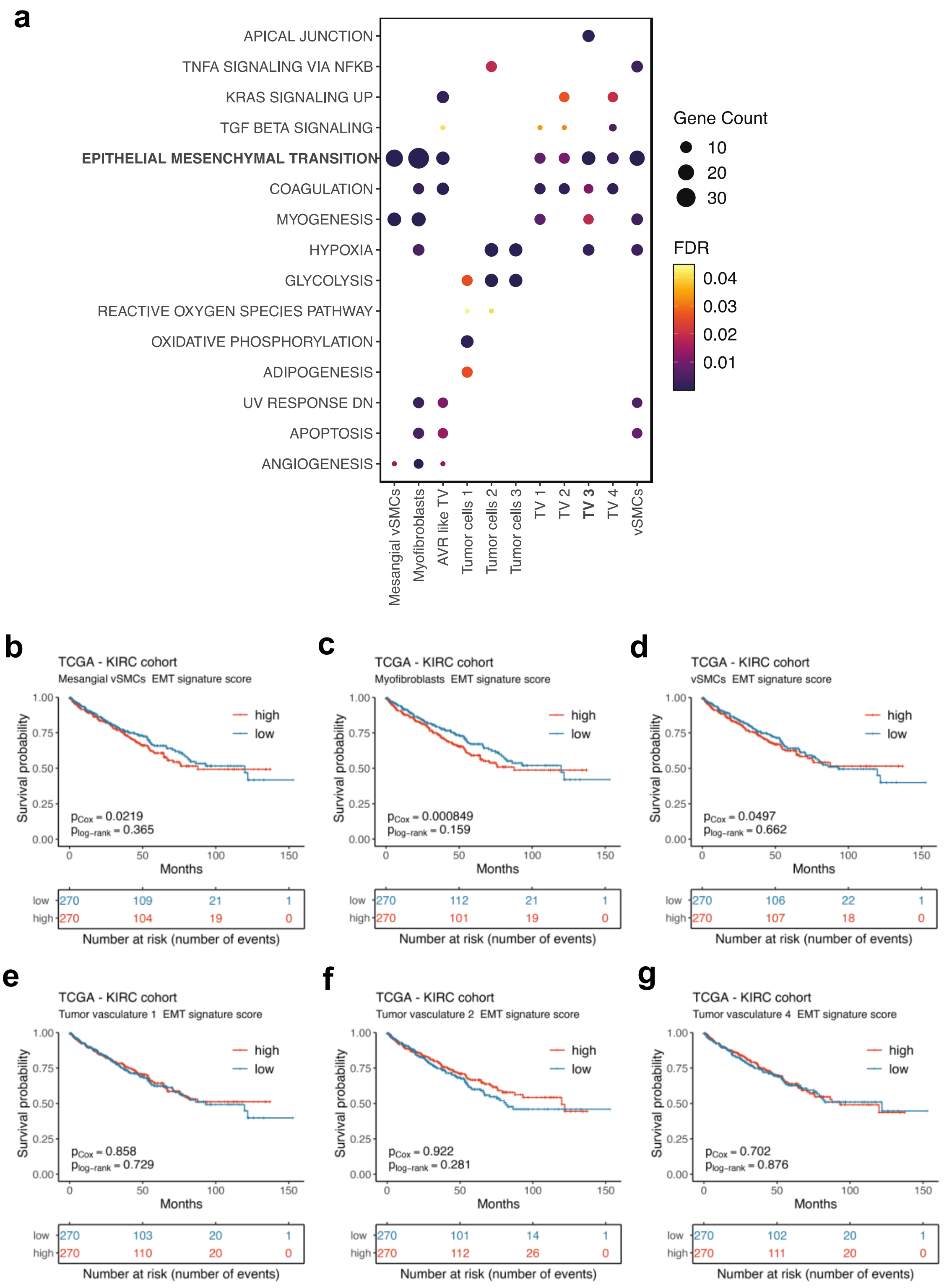


**Supplementary figure S4.** MSigDB Hallmark pathway overrepresentation analysis. **a)** Cell type signature over-representation analysis reveals enrichment of epithelial-mesenchymal transition (EMT) in tumor vasculature and stromal cell populations. **b-g)** None of the stromal and tumor vasculature signature genes overlapping with EMT correlate with overall survival in the TCGA KIRC cohort (except for tumor AVR-like vasculature and tip-like tumor vasculature 3, as shown in Figure 4b-c).

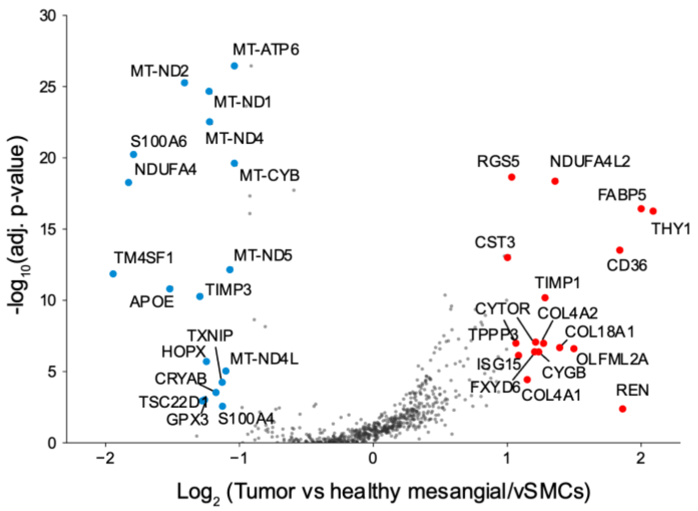


**Supplementary figure S5.** Volcano plot of differentially expressed genes between tumor originating and healthy-adjacent originating cells in mesangial/vSMC population. Genes with fold-change of 2 and adjusted p-value <0.05 are highlighted and considered significant. The asymmetry of the central position of the volcano plot is due to difference in cell size with tumor cells having higher fraction of non-zero genes. This effect was not corrected to maintain consistency in differential gene expression analysis performed.

**Table 1.** Cluster annotation by top 25 DEGs.

| Cluster No in Supp. Table S2 | Annotation | Top 25 genes and references |
|---|---|---|
| 0 | Tumor vasculature 1 | PLVAP[1-3], SPRY1, SPARC, PLPP1, COL4A1[4], VWA1, SPARCL1[5,6], VWF[2,4,7], HSPG2[2], PLPP3[4], GNG11, COL4A2[4], RGCC, RBP7, IGFBP7[4], TIMP3[8], RAMP2[8], IFI27, EGFL7[9], FLT1[1,2], RAMP3, ESAM, INSR[4], GSN, A2M |
| 1 | Tumor vasculature 2 | IGFBP3[10], ENPP2[7], RBP7, ESM1, SPARCL1[5,6], FLT1[1,2], A2M, PLVAP[1-3], SPRY1, STC1, ANGPT2, INSR[4], PRSS23, PLPP3[4], VWF[2,4,7], PLPP1, GNG11, PECAM1[1,2,8], SPARC, EFNB2[11], GJA1, TIMP3[8], IGFBP7[4], IFI27, EPAS1[12] |
| 2 | Proximal tubule | MIOX[8,13], ALDOB[8], MT1G[8,13], GATM[8,13], GSTA2[8], GSTA1[8], ASS1[8,14], FABP1[8], NAT8[8], GPX3[8,14], PDZK1IP1[2,8,13], PCK1[8], MT1H[8], RBP4[8], DCXR[8], HPD[8], SUCLG1[8], ECHS1[8], FBP1[8], UGT2B7[8], BBOX1[8], GLYAT[8], RIDA[8], PEPD[8], CYB5A[8] |
| 3 | Plasma cells | IGHA1[15,16], IGLC3[16], IGHM[16], IGLC2[15,16], JCHAIN[16], IGKC[16], MZB1[15,16], SSR4, DERL3, SEC11C, FKBP11, CD79A[15,16], HERPUD1, PRDX4, HSP90B1, XBP1, FKBP2, ITM2C, DNAAF1, CD27, SELENOS, ATF4, LMAN2, SDF2L1, SSR3 |
| 4 | TAM 1 | CCL3L1, CCL3[2,17,18], IL1B[2,19,20], CXCL8[17,20,21], IER3, CCL4L2, TNF[19,20], CXCL3[17], CXCL2[17], MS4A7, BCL2A1, C1QC[22], PLAUR, C1QA[20,22,23], HLA-DRA[20], GPR183, EIF4E, C1QB[20,22], CD83, NFKBIA, HLA-DPA1[20], HLA-DPB1[20], CCL20[2], CD74[20,23], HLA-DQB1[20] |
| 5 | Principal cells | FXYD4[11,13], AQP2[8,11,24], AQP3[8,11,24], GDF15, WFDC2, EPCAM, HSD11B2[11,13], KRT19[13], TACSTD2, CLU[25], CDH16, SLPI[13], AIF1L, ELF3, DEFB1[14], CLDN4[8,26], ATP1B1, RASD1, FOLR1, S100A2, COBLL1, GSTM3, RALBP1, SPINT2, CD24 |
| 6 | vSMCs | TAGLN[11,13,27], ACTA2[11,13,27], MYL9, ADIRF, TPM2, RGS5, CALD1, C11orf96, PLN, REN, MYH11[11,13], MGP, RERGL[11], BGN, PPP1R14A, PLAC9, CSRP2, SOD3, DSTN, RGS16, NOTCH3[27], IGFBP7, MFGE8, MYLK, CCL2 |
| 7 | Glomerular endothelium | IGFBP5[4,8,28], EMCN[8,11,13], CRHBP[2,8], PLAT[8,11], SOST[2,8], SLC9A3R2[8,28], EHD3[8,11], MGP[8], PLPP3[8], RNASE1[8], IFI27[8], TGFBR2[8], FCN3[8], PLPP1[8], ID1[8], RAMP2[8], GNG11[8], IFITM3[8], ADGRF5[8], PTPRB[8], SLC14A1[8], EPAS1[8], MEIS2[8], TIMP3[8], APP[8] |
| 8 | Classical monocytes (CD14+) | S100A9[15,20,29], S100A8[15,20,29], LYZ[29], G0S2, FCN1[15], IL1B, EREG, THBS1, CXCL8, S100A12[15,29], VCAN[15,29], PLAUR, CXCL2, BCL2A1, CTSS, AREG, C5AR1, NAMPT, PPIF, LST1, NLRP3, IER3, SOD2, CEBPB, ATP2B1-AS1 |
| 9 | Mast cells | TPSB2[8], TPSAB1[8,20], CPA3[8], AREG[8], CD69[8], CTSG[8], MS4A2[8], LMNA[8], LTC4S[8], HPGDS[8], RGS13[8], RHEX, RGS2, KIT[20], HPGD, VWA5A, SRGN, FOSB, CPM, GATA2, NFKBIA, GPR65, ANXA1, LMO4, PPP1R15A |
| 10 | Distal convoluted tubule/connecting tubule | DEFB1[11,14], CALB1[11,13,30], HSD11B2, KLK1[11], KNG1, ATP1B1, TMEM52B[13], CA12, MAL, WFDC2[8], SLC12A3[11,13,30], RHCG[8], GDF15, NFE2L2[8], EPCAM, CDH16, HMGCS2, FXYD2[11,14], SERPINA5, CA2, KCNJ1[1,31], KRT19, MTRNR2L8, EMX1, CLDN4[8] |

| | | |
|---|---|---|
| 11 | Mesangial/vSMCs | RGS5, BGN[25,32,33], REN[34], PLAC9, ADIRF, IGFBP7, CALD1[33], MGP[13], MYL9[33], NDUFA4L2, HIGD1B, TPM2, ID3, FABP4, NOTCH3, CD36[35], PDGFRB[11,13,27], C11orf96, CSRP2, FRZB, TAGLN[11], TPPP3, THY1[30], COL1A2, LHFPL6 |
| 12 | Mast cells | TPSB2[8], TPSAB1[8,20], CPA3[8], CTSG[8], CD69[8], AREG[8], HPGDS[8], RHEX, MS4A2[8], HPGD, VWA5A, CCL2, RGS13[8], RGS2, LTC4S, SRGN, ANXA1, LMNA, GPR65, KIT[20], ALOX5AP, PPP1R15A, LAPTM4A, FCER1A, AC020571.1 |
| 13 | Cycling | HIST1H4C, STMN1[8], HMGB2[36], PCLAF, UBE2C[8,36], HMGN2, TUBB, CKS1B, MKI67[8,23,36], TUBA1B, H2AFV, H2AFZ, CENPF[36], PTTG1, TYMS[8,36], RRM2[36], HMGB1, PCNA[36], DUT, CLSPN[36], NUSAP1[36], GZMK, GZMA, CXCL13, DEK |
| 14 | IGHG-high plasma cells | IGKC[16], IGHG3, IGHG1, IGLC2[15,16], IGHG4, IGLC3[16], JCHAIN[16], MZB1[15,16], IGHGP, IGHG2, SSR4, SEC11C, XBP1, CD79A[15,16], FKBP11, DERL3, HSP90B1, ITM2C, PRDX4, FAM30A, ERLEC1, SPCS1, IGHM, DNAAF1, KLF13 |
| 15 | Non-classical monocytes (CD16+) | LST1[37], SMIM25, FCN1[15], AIF1[37], G0S2, LYPD2[38], COTL1, SAT1, FCER1G, CTSS[37], BCL2A1, IL1B, LILRB2[15], PLAUR, FCGR3A[20,37,39], C5AR1, TYROBP, MS4A7, TIMP1, NAP1L1, POU2F2, IFI30, STXBP2, CALHM6, NEAT1 |
| 16 | Descending vasa recta | TIMP3[8], CLDN5[2,13], AQP1[2,11,40], SLC9A3R2[28], TM4SF1, IFI27, RAMP2[8], RBP7, SLC14A1[11,13,40], DEPP1, ID1[2], CRIP2, SERPINE2[2], PPP1R14A, SRP14, PALMD[13], FAM107A, PLPP1, ICAM2, RNASE1, SSUH2, KCTD12, S100A6, ABI3BP, IFITM3[8] |
| 17 | Proximal tubule | MT1G[8,13], ALDOB[8], MT1H[8], GPX3[8,14], FABP1[8], PDZK1IP1[2,8,13], MIOX[8,13], HPD[8], MT1X[8], CXCL14[8], ASS1[8,14], GATM[8], NAT8[8], MT1F[8], GSTA1[8], SUCLG1[8], SMIM24[8], FXYD2[8,13], RBP5[8], RIDA[8], MT1E[8], SPP1[8], UGT2B7[8], LGALS2[8], ALB |
| 18 | Tumor vasculature 3 | LY6H, CLDN5[2,13], PGF, COL4A1[4], APOD, SPARC[41], COL4A2[4], HSPG2[2], CHST1, TNFRSF4, PXDN, VWF[2,4,7], LAMA4, GNG11, CCDC85B, PECAM1[1,2,8], SPARCL1[2], PRSS2, FSCN1[4], SPRY1, CRIP2, LXN, ICAM2, TCF4, CD93 |
| 19 | Tumor cells 1 | FABP7[42], GC, CD24[43,44], ANXA4, SLC17A3[2,23], NDRG1[45], UBD, PLIN2[46], CRYAB[23], CLU[23], SPATS2L, FHL1, MDK, HLA-G, MGST2, GSTA1, HMGN3, CISD1, ATP1B1, NNMT[2], SLC13A1, CXCL14, SOD2[23], HSPB8, VCAM1[23] |
| 20 | Outer medulla type A intercalated cells | SPINK1[8], TMEM213[8,13], ATP6V1G3[11,36,47], SLC4A1[1,11,13], DEFB1, ATP6V0D2[11,13,36], RTN4[8], CKB[8], RHCG[8], EPCAM, SMIM6, ATP6AP2[8,13], MAL[8], FAM24B[8], CA12[8], LGALS3, HSD11B2, C12orf75[13], ADGRF5[11,13,47], ATP6V0B[8], BSG, DHRS7, ATP6V0A4, CLCNKB[1,11], ATP1B1<br>Also positive for marker SLC26A7[48] |
| 21 | Myofibroblasts | COL1A1[49], COL1A2[49], TIMP1[49], MGP[8], TAGLN[8], COL3A1[49], ACTA2[1,8,49], BGN[8,10], FN1, TPM2[8], DCN[10], CALD1[8,10], MYL9[8], IGFBP7[8], POSTN, RGS5[8], LUM[49], COL6A2[8], SPARC[8,10], PLAC9[8], COL4A2, AEBP1, COL6A1[10], PPP1R14A, COL4A1 |

| | | |
|---|---|---|
| 22 | Tumor cells 2 | NNMT[2], NDUFA4L2[2,23,50], CD24[43,44], VEGFA[23], PDK4, CRYAB[23], ANGPTL4[2,51], PLIN2[46], CLU[23], TMEM176B, NDRG1[45], TMEM176A, RARRES2, HILPDA, BNIP3[52], ANXA4, RNASET2, CYB5A, CCDC146, KRT8, CCND1[52], ENO1, CXCL14, KRT18, EGR1 |
| 23 | Thick ascending limb of LOH | DEFB1[8,11], UMOD[11,14,30], PCP4[8], FXYD2[8,14,31], KNG1[8,13], CKB[8], GSTM3[8], MPC1[8], LDHB[8], PEBP1[8], MRPS6[8], PPP1R1A[8], CYSTM1, BEX3, SLC25A5[8], SLC12A1[8,11,14], CBR1, ATP5F1A[8], SLC25A4[8], CD24[8], TCIM, ATP5MC3[8], UQCRFS1[8], PRDX2[8], MAL |
| 24 | Tumor AVR-like vasculature | ACKR1[1,2,8], VWF[2,4,7], RNASE1[8], CLU[2], IFI27[8], TFF3, CD59, HYAL2[8], RAMP3[8], FKBP1A, FN1, TM4SF1[8], IFITM3[8], PTGDS, PECAM1[1,2,8], ADIRF, DNASE1L3[2,13], S100A6, IGFBP4[8], SLCO2A1[13], ECSCR, FAM167B, RAMP2[8], IL33, TGM2 |
| 25 | Regulatory T cells | LTB, TNFRSF18[23,53], IL32, BATF[53], TIGIT[53,54], CARD16, CORO1B, TNFRSF4[53,54], TRAC[23], TRBC1, S100A4, LINC01943, FOXP3[23,53,54], CD3D[41,53], LAIR2, AC133644.2, LINC02195, CD27, CTLA4[53,54], PMAIP1, TBC1D4, DUSP4, CYTIP, SPOCK2, ICA1 |
| 26 | Thick ascending limb of LOH | UMOD[11,14,30], DEFB1[8,11], KNG1[8,13], SLC12A1[8,11,14], ATP1A1, ATP1B1, S100A2, CLDN10, SFRP1, KCNJ1[8,11], CA12, MAL, CKB[8], TMEM52B, CDH16, UCHL1, GSTM3, PCP4[8], SERPINA5, CLCNKB, TSPAN8, MPC1[8], SPP1, MTRNR2L8, EGF |
| 27 | Tumor cells 3 | FABP7[42], NDUFA4L2[2,23,50], CD24[43,44], NNMT[2], BNIP3[52], HILPDA, CRYAB[23], PLIN2[46], LDHA[52], RARRES2, ANGPTL4[2,51], ANXA4, GAPDH, ENO1, TPM1, NDRG1[45], CLU[23], TPI1, VDAC1, SERPINA1, TMEM176A, NUPR1, KRT18, VEGFA[23], EGLN3[55] |
| 28 | Tumor vasculature 4 | PLVAP[1-3], GNG11, SPARC, RBP7, SPRY1, HSPG2[2], VWF[2,4,7], IFI27, RAMP2[8], PLPP1, COL4A1[8], SPARCL1[5,6], IGFBP7[4], TIMP3[8], MTRNR2L8, EGFL7[9], INSR[4], GSN, RAMP3, ESM1, VWA1, IFITM3[8], ECSCR, TCF4, SLC9A3R2 |
| 29 | Podocytes | PTGDS[8], DCN[8], NPHS2[8,11,13], EMCN, IGFBP5, TNNT2[8], IGFBP2[8], HTRA1[8], PCOLCE2[8], CTGF, PLAT, PODXL[8,13,56], TPPP3[8], MYL9[8], CRHBP, APOD[8], MME[8], HPGD, CLIC5[8,13,56], TCF21[8,56], CDKN1C[11,47], SLC9A3R2, ID1, BST2[8], AIF1[8] |
| 30 | TAM 2 | APOC1[39], APOE[20,23,39], C1QB[20,22], C1QA[20,22,23], C1QC[22], CTSD[39], HLA-DRA[20], TYROBP, TREM2[54], HLA-DPA1[20], HLA-DQA1[20], HLA-DRB1[20], FCER1G, PSAP, HLA-DPB1[20], LYZ, HLA-DQB1[20], CTSB, GPNMB[23,39], NPC2, CST3, CD68[1,2,54], CD74[20,23], HLA-DRB5[20], CFD |
| 31 | CD8 T cells | DUSP4[41], GZMK[41,53], CST7, LYST, CCL5[53], TRBC2, CD8B[53], CD3D[41,53], RGS1, CD27[41], CREM[41], RGS2[41], NKG7[53], TRAC[53], TNFRSF9[41,53], CD8A[41,53], TRBC1, HSP90AA1[41], GZMA[23], SRSF7, CD2, PMAIP1, CMC1, RUNX3[57], SYTL3 |
| 32 | Type A intercalated cells | ATP6V1G3[11,36,47], DEFB1, TMEM213[8,13], C12orf75[8], SPINK1[8], CKB[8], FXYD2[8], MAL[8], ATP6V0D2[11,13,36], FAM24B[8], SMIM6, NUPR2, LGALS3[8], CYSTM1[8], CA12[8], CLCNKB[1,11], RTN4[8], SLC25A5[8], SLC25A39[8], BSG[8], |

| | | |
|---|---|---|
| | | HOXB-AS3, ERP27[13], COX7A1, SLC4A1[1,11,13], ADGRF5[11,13,47] |
| 33 | B cells | CD79A[15,16,23], CD37[16], IGHM, IGKC, MS4A1[15,16], LTB[16], BANK1, IGHD, LINC01781, LINC00926, CD83, RALGPS2, FAM30A, CD52, BIRC3, EEF1B2, RPS5, POU2F2, SPIB, RPL32, RPL18A, LINC01857, RPS8, CD79B[15,16], CD55 |
| 34 | Thin ascending limb of LOH | WFDC2, S100A2[58], SLPI, SOD3, PAPPA2[59], MMP7, DEFB1, ITM2C, TACSTD2[13], MAL, SLC12A1[13], PCSK1N, IGFBP6, CLDN10[13,58], S100A6[58], UMOD[13,58], CLU[14], TSPAN8[13], CA12, MUC1, CD24, CLDN3, ATP1A1, KRT7, EPCAM |
| 35 | Ascending vasa recta | DNASE1L3[2,13], IGFBP5[28], RNASE1[8], RAMP3[8], EMCN[8,13], CAVIN2[8], IFI27[8], ID1, TMEM88, RAMP2[8], SLC9A3R2[28], CA4, GNG11[8], IFITM3[8], FAM167B, ENG[8], TIMP3[8], TM4SF1[8], PLPP3, HYAL2[8], PCAT19, IGFBP4[8], PLAT[8], MEIS2[8], GIMAP7 |
| 36 | Type B intercalated cells | KRT7[8], WFDC2[8], ATP6V1G3[11,36,47], TMEM213[8,13], S100A2[8], CD9[8], CA12[8], MTRNR2L12, DEFB1, TSPAN8[8], MAL[8], RARRES2[8], CDA[8], ATP6V0B[8], ATP6V0D2[8,11,13], ATP6AP2[8], LGALS3[8], ATP6V0A4[8], ATP6V1B1[8,11], MTRNR2L10, SMIM24[8], EPCAM, CKB, MTRNR2L1, MTRNR2L8<br>Also positive for markers: SLC26A4[11,13,60], HMX2[11], SPINK8[11] |
| 37 | TAM 3 | HLA-DPB1[20], HLA-DPA1[20], APOC1[39], CST3, HLA-DRA[20], TYROBP, AIF1, HLA-DQA1[20], C1QA[20,22,23], HLA-DQB1[20], C1QB[20,22], CD74[20,23], MS4A6A, C1orf162, FTL, LYZ, RGS10, NPC2, CD68[1,2,54], LST1, C1QC[22], SAT1, IER3, HLA-DMA[20], HLA-DRB1[20] |
| 38 | Epithelial progenitor-like cells | CRYAB[8], SLPI[8], MMP7[8], CLU[8,61], CLDN4[8], WFDC2[8], TACSTD2[8], CD24[8,58,61], CTGF, CLDN3[8], NUPR1, MT1E, ELF3[8], EGOT, TPM1[8,58], KRT19, MAL[8], PIGR, S100A13[8], SPINT2[8], SOX4[61], RASD1, ATF3, MT1X, BCAM[8] |
| 39 | Cytotoxic T cells | XCL1[53], XCL2[53], KLRB1[62], TRDC, KLRC1, KRT86, CD7, KLRD1[62], HOPX[53], GNLY[53,62], CCL4[53], KRT81, CCL5[53], ZNF683[53], LINC01871, CTSW[62], HCST, IL2RB, AREG, ZFP36L2, BTG1, REL, FAM177A1, MATK, CMC1 |
| 40 | TAM 4 | C1QB[20,22], C1QA[20,22,23], C1QC[22], SELENOP[20], FOLR2[20,23], RNASE1, HLA-DPA1[20], CD14, HLA-DRA[20], MS4A6A, MS4A7, HLA-DRB1[20], AIF1, HMOX1, MS4A4A[20], TYROBP, HLA-DPB1[20], NPC2, CD74[20,23], HLA-DQA1[20], CST3, HLA-DQB1[20], FCGRT, SLC40A1[21,39], FCER1G, PLTP[20], BLVRB |
| 41 | NK cells | GNLY[63], GZMB[63], FGFBP2, NKG7[63], KLRD1[63], KLRB1[53], PRF1[63], GZMH[63], SPON2[63], PLAC8, CLIC3, KLRF1[63], CD247[63], AREG, GZMM, HOPX, CST7[63], TRDC, CMC1[63], ARL4C, CTSW, CCL4[63], PTGDS[63], GZMA[63], EFHD2 |
| 42 | Resting/memory T cells | IL7R[23,53], CD52[53], LTB[64], KLRB1[53], ZFP36L2[41], RPS29, CD48, CXCR4[41], LYAR, BTG1, ANXA1, STK4, RPS27, EML4, PPP2R5C, RPS3, ACAP1, RPS15A, SPOCK2, ISG20, FYN, CD3D[41,53], PDE4B, CD2, RPLP2 |
